# Longitudinal white matter changes associated with cognitive training

**DOI:** 10.1101/2020.11.24.396119

**Authors:** E.S. Nichols, J. Erez, B. Stojanoski, K.M. Lyons, S.T. Witt, C.A. Mace, S. Khalid, A.M. Owen

## Abstract

Improvements in behaviour are known to be accompanied by both structural and functional changes in the brain. However whether those changes lead to more general improvements, beyond the behaviour being trained, remains a contentious issue. We investigated whether training on one of two cognitive tasks would lead to either near transfer (that is, improvements on a quantifiably similar task) or far transfer (that is, improvements on a quantifiably different task), and furthermore, if such changes did occur, what the underlying neural mechanisms might be. Participants trained on either a verbal inhibitory control task or a visuospatial working memory task for four weeks, over the course of which they received five diffusion tensor imaging scans. Two additional tasks, a test of verbal reasoning and a test of spatial span, served as measures of near transfer for the inhibitory control task and spatial working memory task, respectively. These two tasks also served as measures of far transfer for the alternate training task. Behaviourally, participants improved on the task that they trained on, but did not improve on cognitively similar tests (near transfer), nor cognitively dissimilar tests (far transfer). Extensive changes to white matter microstructure were observed, with verbal inhibitory control training leading to changes in a left-lateralized network of fronto-temporal and occipito-frontal tracts, and visuospatial working memory training leading to changes in right-lateralized fronto-parietal tracts. Very little overlap was observed in changes between the two training groups. On the basis of these results, we suggest that near and far transfer were not observed because the changes in white matter tracts associated with training on each task are almost entirely non-overlapping with, and therefore afford no advantages for, the untrained tasks.

## 1 Introduction

It is well documented that improvements in behaviour, particularly motor behaviour, are accompanied by both structural and functional changes in the brain. For example, learning to juggle is associated with white matter microstructure changes in visuomotor regions (Scholz et al., 2009), while levels of functional connectivity observed while practicing a finger-tapping motor sequence can predict the amount of learning in a later session (Bassett et al., 2011). The same is true of cognitive learning; London (UK) taxi drivers, who have achieved a high level of expertise in spatially navigating the city, have greater hippocampal grey matter volume than controls (Woollett & Maguire, 2011), while studying for the Law School Admission Test leads to white matter changes within the frontoparietal network (Mackey et al. 2015). Structural and functional changes after learning a cognitive task in the laboratory have also widely been observed (e.g. Lampit et al., 2015; Thompson et al., 2016).

One interesting observation arising from this literature is that training on a cognitive task typically results in improvements on that task, but not on others, even when they are quite closely related. As an example, in well-known series of studies involving taxi drivers (Maguire et al., 2006; Woollett & Maguire, 2011), the drivers were better at navigating through London, although they were generally no better at other cognitive tasks. In fact, they performed worse than controls in acquiring new visuospatial information (Maguire et al., 2006). This speaks to one of the most contentious issues in the brain training literature; that is, whether training on cognitive tasks can lead to generalized improvements in cognition. While some authors have suggested that training on one cognitive task leads to improvements on unrelated tasks (Au et al., 2015; Jaeggi et al., 2008), a considerable number of studies have failed to show such effects (Owen et al., 2010; Stojanoski et al., 2018, 2020). Indeed, while the holy grail of the brain training literature is so-called ‘far transfer’ (i.e., where training on one task improves performance on a completely unrelated second task), many studies have failed to even demonstrate ‘near transfer’ (i.e., when training on one task improves performance on a similar, cognitively related second task) (Simons et al., 2016). Whether brain training “works” or not will likely remain a contentious issue. Regardless, what is lacking from the literature is any clear neuroscientific explanation for *how it would work, assuming it works at all*.

If we first take the position that brain training, in general, does not lead to any transfer, we can generate a number of hypotheses for why that may be the case. Unlike many aspects of motor control, cognitive processes rarely have a one-to-one mapping between the behavioural tasks used to test them and structures in the brain. For example, there is no “digit span” area of the brain comparable to the brain regions that are known to initiate a simple motor function such as finger tapping (Cramer et al., 1999). Cognitive processes generally recruit extensive networks of brain regions, none of which are uniquely and singularly devoted to performing a specific cognitive task (Crittenden et al., 2016; Crittenden & Duncan, 2014; Duncan, 2010; Duncan & Owen, 2000). Even though training on a cognitive task evokes changes at the network level (Bassett et al., 2011; Finc et al., 2020), it is possible that transfer does not occur between tasks because the networks that drive those tasks are not similar enough for changes in one to improve the other. To illustrate this point, consider the relationship between spatial span and digit span. Although on the surface they appear to be similar tasks, they differ in both obvious and less obvious ways. Clearly, they differ in terms of modality of the memoranda; however, it is also known that the strategies that are adopted to solve these tasks are quite different (Bor et al., 2003, 2004). While the most common strategy that improves performance on digit span is chunking, where several numbers are grouped together into single units to be remembered, spatial span relies more heavily on pattern recognition (for discussion, see Owen, 1997). These differences may outweigh any surface similarities that these two tasks may have (e.g. both tasks rely on storing and repeating ordered information).

Let us now take the alternate position that brain training on one task does lead to benefits on other tasks, and consider what the neuroscientific mechanism for that might be. Presumably, the two tasks (the trained and the untrained) must recruit networks that are similar enough that strengthening one is sufficient to lead to performance enhancements on the other. This leads to two testable predictions: first, if training on a task leads to transfer, those two tasks should recruit largely overlapping brain networks. Second, for quantitatively similar cognitive tasks, a significant amount of network overlap and a correspondingly large amount of transfer should be observed. In contrast, cognitively dissimilar tasks should involve less network overlap, and thus, less transfer.

An important step in testing these predictions is not to choose tasks based on an intuitive sense that they are similar or different (e.g., see Digit Span and Spatial Span example above). Fortunately, it is possible to select tasks based on quantifiable measures of similarity between them, thereby operationalizing how transfer is defined and measured. For example, based on a factor analysis computed by Hampshire et al. (2012) that grouped 12 cognitive tasks into three functionally and anatomically distinct neural components, the assigned factor and the corresponding factor loadings of each cognitive task can be used to guide the choice of training and transfer tasks. Using that method, we selected a test of inhibitory control as a training task for the current study, and a test of grammatical reasoning to assess near transfer because, among a group of 44,000 participants, they loaded heavily on the same factor (Hampshire et al., 2012). Similarly, a test of spatial working memory was selected as a second training task (in a separate group of individuals), and a test of spatial span was used as a second test of near transfer because, in the same group of 44,000 participants, they loaded heavily on the same factor as one another. Importantly, these two sets of tests loaded on entirely different factors and were functionally and anatomically dissociable, confirming that there is very little overlap between the demands of each. Therefore, the two near transfer tasks (Grammatical Reasoning and Spatial Span) also serve as ideal measures of far transfer for the spatial working memory task and inhibitory control task, respectively.

Based on previous work (e.g. Caeyenberghs et al., 2016; Lampit et al., 2015; Thompson et al., 2016), we hypothesized that training on a cognitive task will lead to behavioural improvements on that task that are associated with specific structural changes in the brain’s white matter microstructure. Second, based on our own previous work (Owen et al., 2010; Stojanoski et al., 2018, 2020) we hypothesized that training on a cognitive task will not lead to improvements on a second cognitive task, regardless of whether it is quantifiably similar (near-transfer) or quantifiably different (far-transfer). Finally, we hypothesized that this lack of transfer effect is underpinned by the different structural changes that are associated with learning the two training tasks. Specifically, we expected that improvements in spatial working memory with training would lead to structural changes that are non-overlapping with improvements observed with inhibitory control training.

## 2 Method

### 2.1 Participants

Twenty-one participants (19 females) ranging in age from 18-36 (*M* = 25.2, *SD* = 5.09) were enrolled in the study. Participants were recruited via posters on bulletin boards around Western University campus, London, Canada, and received monetary compensation for their participation. All participants gave written informed consent prior to study commencement. The study was approved by Western’s Research Ethics Board (#108475) and performed according to the ethical standards laid down in the 1964 Helsinki Declaration and its later amendments. Due to scanner issues, four participants were not able to complete the final two follow-up scans and were excluded from further analysis. One additional participant withdrew after the initial scanning session and thus was also removed from further analysis. A final sample size of 16 participants was analyzed.

### 2.2 Training Procedure

Participants were randomly assigned to train on either ‘Double Trouble’, a modified Stroop test of inhibitory control described below, or ‘Self-Ordered Search’, a visuo-spatial working memory task, also described below. They completed three to five at-home training sessions per week and five in-person scans, and a schematic of the training protocol is provided in Figure 1. Both resting state and task-based fMRI data were also collected, but were not analyzed for the current study. During the first scanning session, the task instructions were explained to participants and any questions were answered to ensure full understanding. Additionally, text and visual instructions were presented at the beginning of each training session as a reminder.

**Figure 1.**
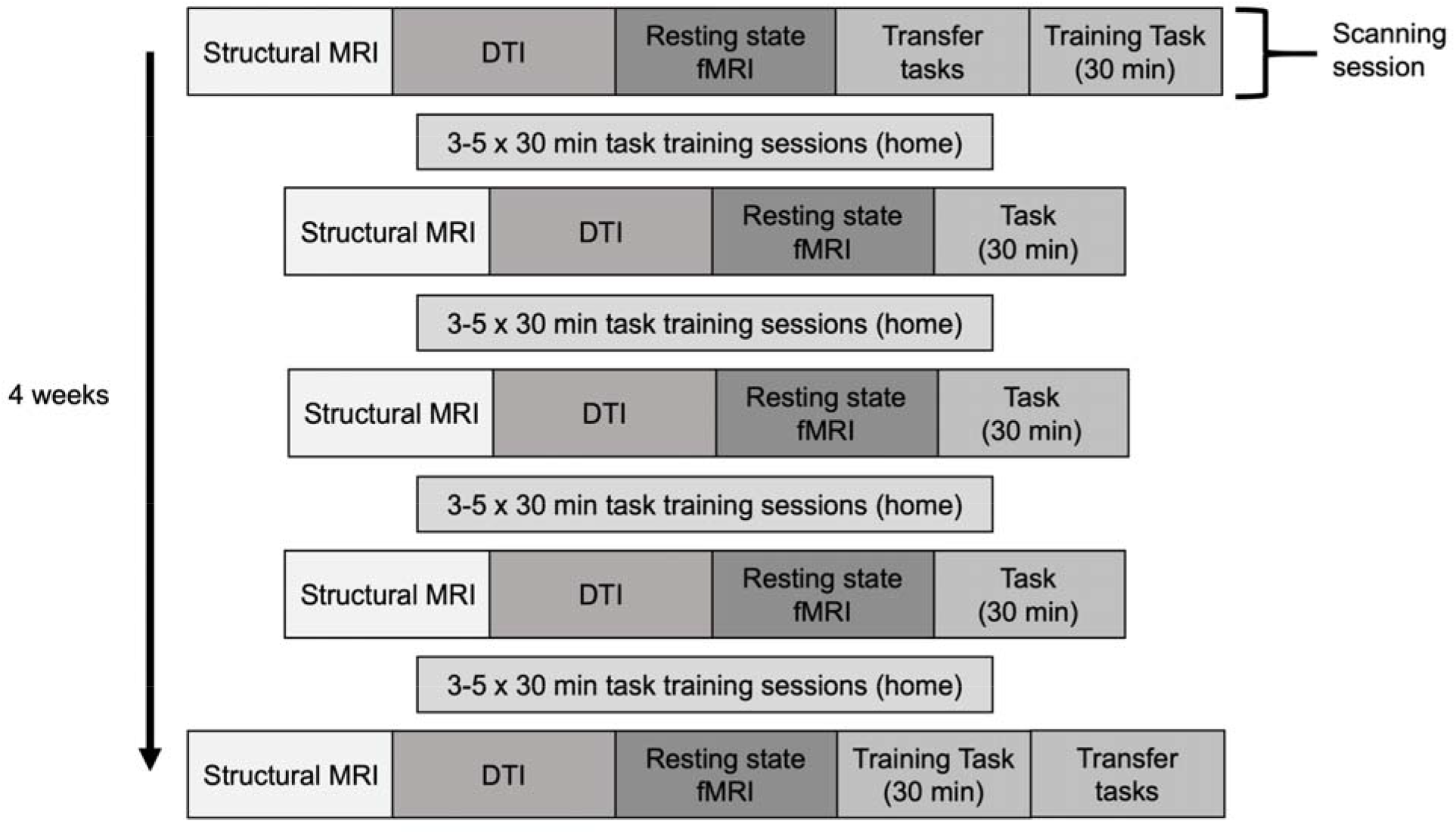
Study Procedure. Five scanning sessions were completed over the course of 4 weeks, with one week in between each session. Participants trained on the task at home between scans.

#### 2.2.1 At-home training

All participants completed a minimum of three and a maximum of five training sessions per week over the course of four weeks, with at least 24 hours between each session, but no more than 72 hours without training. Each training session took approximately 30 minutes and was completed at home using the Cambridge Brain Sciences online platform at cambridgebrainsciences.com (Hampshire et al., 2012).

#### 2.2.2 In-scanner training

Participants completed five imaging sessions throughout their cognitive training to track changes in their white matter microstructure. The first imaging session was completed prior to the start of their cognitive training. The subsequent four imaging sessions occurred with the at-home training sessions between each. During all imaging sessions participants underwent a structural MRI, DTI, and resting-state fMRI scan before completing their training task for approximately 30 minutes. During the first and last scanning sessions, participants also completed two tasks used to measure cognitive transfer, described below. To eliminate practice effects, participants were allowed to practice the transfer tasks twice each, and the score on their third attempt was taken as their measurement of performance on that test.

### 2.3 Tasks

Four computerized cognitive tasks from an online cognitive testing battery (cambridgebrainsciences.com) were used in this study. The two training tasks, Double Trouble and Self-Ordered Search, were chosen as they tax different cognitive domains, specifically inhibitory control and verbal abilities, and visuo-spatial processing/working memory, respectively (see Hampshire et al., 2012). Using tasks that tap into functionally and anatomically dissociable cognitive domains allowed us to determine whether functional and structural changes following cognitive training are specific to the training task, are more domain-general changes, or are some combination of both.

Two additional tasks were used as measures of cognitive transfer. The first task was the ‘Grammatical Reasoning’ task from the Cambridge Brain Sciences battery, which was used to assess near-transfer for the Double Trouble training group and far transfer for the Self-Ordered Search group. In a factor analysis of 44,000 participants, Grammatical Reasoning loaded heavily on the same factor as Double Trouble (factor loadings = 0.66 and 0.51, respectively) and, through fMRI was shown to recruit a similar functional network in the brain (Hampshire et al., 2012). In contrast, the Self-Ordered Search task did not load heavily on that factor (factor loading = 0.16) and recruited a different functional network. The second task, ‘Spatial Span’, is another test of spatial working memory and was used to assess near transfer for the Self-Ordered Search group and far transfer for the Double Trouble group. In the same factor analysis, Spatial Span loaded heavily on the same factor as Self-Ordered Search (factor loadings = 0.69 and 0.62, respectively), and the Double Trouble task did not load heavily on that factor (factor loading = 0.22). These tasks were completed in-scanner prior to completing the training task during the first scanning session, and after completing the training task during the last scanning session.

#### 2.3.1 Double Trouble

The Double Trouble task is a modified version of the classical Stroop test (Stroop, 1935) and measures inhibitory control, verbal ability, and attention. On each trial, a probe word “red” or “blue” is displayed at the top of the screen in either red or blue font. The participant must select the word at the bottom of the screen that describes the font colour of the probe word, inhibiting any response based on the probe word (“red” or “blue”) itself. The two response choices are the words “red” or “blue” and are also displayed in either red or blue font, which ‘doubles’ the inhibitory load because participants have to inhibit any response based on font colour. The word and the colour of the font may be congruent or incongruent for both the probe and the answer choices. The participant is given 90 seconds to complete as many trials as possible. Their score increases by one point each time they make a correct response and decreases by one point each time they make an incorrect response. Participants assigned to this condition completed approximately 20 90-second runs within each 30-minute training period.

#### 2.3.2 Self-Ordered Search

The Self-Ordered Search task assesses visuospatial working memory (Owen et al., 1990). A set of squares in random positions within an invisible five by five grid is displayed on the screen. Participants click on squares, which ‘open’ to reveal whether there is a “token” inside. After a token is found, it is hidden within another square, and the participant must locate it again. Within any trial, a square will never be used to hide a token more than once and the number of tokens that must be found in each trial equals the number of squares in that trial. Participants must avoid squares in which they have already discovered a token and squares they have already searched whilst looking for the current token. If they re-click a previously discovered or searched square the trial ends and the next trial begins with one less square in the grid. If they find all tokens without making an error, a new trial begins with one extra square in the grid. The round begins with four squares and ends after three errors have been made. Their final score is equal to the maximum level they achieved. As this task is not timed, the number of runs completed during a 30-minute period depended on participant performance.

#### 2.3.3 Grammatical Reasoning

Grammatical Reasoning is based on Alan Baddeley’s three-minute grammatical reasoning test (Baddeley, 1968) and assesses verbal reasoning. On each trial, a written statement regarding two shapes is displayed on the screen, and the participant must indicate whether it correctly describes the shapes pictured below. The participant has 90 seconds to complete as many trials as possible. A correct response increases the total score by one point, and an incorrect response decrease the score by one point.

#### 2.3.4 Spatial Span

Spatial Span is based on the Corsi Block Tapping Task - a tool for measuring spatial short-term memory capacity. Sixteen purple boxes are displayed in a grid. A sequence of randomly selected boxes turn green one at a time (900 ms per green square). Participants must then repeat the sequence by clicking boxes in the same order. Difficulty is varied dynamically: correct responses increase the length of the next sequence by one square, and an incorrect response decreases the sequence length. The test finishes after three errors. The score is the length of the longest sequence successfully remembered.

### 2.4 Behavioural data analysis

Behavioural data were analyzed using R for statistical computing (R Core Team, 2014). Because participants completed multiple rounds of their assigned task during each scanning/training session, the single maximum score was used as that training day’s value. Then, because people trained a different number of times per week, and to accommodate natural minor fluctuations in performance from day to day, we took the maximum score for each participant for training weeks two, three, and four as their overall measure of performance for those weeks, which was then used for data analysis. To assess participants’ trends of learning across their training sessions, scatterplots were created with each week’s highest score plotted over time for each participant. Curve estimation was used to fit linear and logarithmic models to the data to determine the nature of learning trends. Paired-samples t-tests were performed to determine whether logarithmic or linear models fit the learning trend data better for each of the two groups.

To quantify overall task improvement due to cognitive training, and to determine whether there was a significant difference in the amount of learning between the Double Trouble and Self-Ordered Search groups, we conducted a 2 x 2 linear mixed model on scores from Week 1 and Week 5. Scores were first transformed to *z*-scores using population means and standard deviations derived from Wild et al. (2018) to allow for comparison between tests, and the model was built with group (Double Trouble/Self-Ordered Search) and time (Week 1/Week 5) as binary regressors and participants as a random effect.

To assess near transfer, we conducted a linear mixed model on near transfer task scores (that is, Spatial Span scores for the Self-Ordered Search training group, and Grammatical Reasoning scores for the Double Trouble training group). Scores again were first transformed to z-scores using population means and standard deviations (Wild et al., 2018), and the model was constructed with group (Self-Ordered Search/Double Trouble) and time (pre-training/post-training), and participants as a random effect. To assess far transfer, a similar model was constructed but with far transfer task scores (that is, Grammatical Reasoning scores for the Self-Ordered Search training group, and Spatial Span scores for the Double Trouble training group).

### 2.5 Neuroimaging data acquisition

Imaging data were acquired using a 3T Siemens Prisma scanner (Erlangen, Germany) and 32-channel head coil at the Centre for Functional and Metabolic Mapping (Robarts Research Institute, Western University, London, Canada). Whole-brain T1-weighted structural images (repetition time (TR) = 2300 ms, echo time (TE) = 2.98 ms, field of view (FOV) = 256 mm, 256 x 256 matrix, slice thickness = 1 mm, 176 slices) were first obtained. Diffusion-weighted images were acquired in the transverse plane using a single-shot sequence (84 slices with 2 mm slice thickness, voxel size= 2×2mm in-plane, field of view= 210 mm, 137 diffusion directions with b=2000 s/mm2, TR=4 s, TE=59.20 ms; GRAPPA acceleration factor=2).

### 2.6 Neuroimaging data preprocessing

Results included in this manuscript come from preprocessing performed using *fMRIPrep* 1.4.0 (Esteban et al., 2019, 2020) (RRID:SCR_016216), which is based on *Nipype* 1.2.0 (Gorgolewski et al., 2011, 2018) (RRID:SCR_002502).

#### 2.6.1 Anatomical data preprocessing

A total of five T1-weighted (T1w) images were found within the input BIDS dataset. All of them were corrected for intensity non-uniformity (INU) with N4BiasFieldCorrection (Tustison et al., 2010), distributed with ANTs 2.2.0 (Avants et al., 2008) (RRID:SCR_004757). The T1w-reference was then skull-stripped with a *Nipype* implementation of the antsBrainExtraction.sh workflow (from ANTs), using OASIS30ANTs as target template. Brain tissue segmentation of cerebrospinal fluid (CSF), white-matter (WM) and gray-matter (GM) was performed on the brain-extracted T1w using fast (FSL 5.0.9, RRID:SCR_002823) (Zhang et al., 2001). A T1w-reference map was computed after registration of 5 T1w images (after INU-correction) using mri_robust_template (FreeSurfer 6.0.1) (Reuter et al., 2010). Volume-based spatial normalization to two standard spaces (MNI152NLin2009cAsym, MNI152NLin6Asym) was performed through nonlinear registration with antsRegistration (ANTs 2.2.0), using brain-extracted versions of both T1w reference and the T1w template. The following templates were selected for spatial normalization: *ICBM 152 Nonlinear Asymmetrical template version 2009c* (Fonov et al., 2009) [RRID:SCR_008796; TemplateFlow ID: MNI152NLin2009cAsym], *FSL’s MNI ICBM 152 non-linear 6th Generation Asymmetric Average Brain Stereotaxic Registration Model* (Evans et al., 2012) [RRID:SCR_002823; TemplateFlow ID: MNI152NLin6Asym].

Many internal operations of *fMRIPrep* use *Nilearn* 0.5.2 (Abraham et al., 2014) (RRID:SCR_001362), mostly within the functional processing workflow. For more details of the pipeline, see <u>the section corresponding to workflows in *fMRIPrep’*s documentation</u>.

#### 2.6.2 Diffusion-weighted images preprocessing

Diffusion-weighted images were first checked for quality using DTIPrep (Oguz et al., 2014), an automated toolkit. Images were first converted to NRRD file format and checked for header information including correct image dimensions, spacing, and orientation. DTIPrep then ensures correct diffusion gradient orientations and *b-values*. Rician noise removal was performed, followed by artifact detection and removal. Images were then co-registered to an iterative average over all baseline images, followed by eddy-current correction and motion correction, including gradient direction adjustments. A second round of motion detection was the performed to ensure that registration was successful.

### 2.7 DTI analysis

Voxelwise statistical analysis of the fractional anisotropy (FA) data was carried out using TBSS (Tract-Based Spatial Statistics) (Smith et al., 2006), which is part of FSL (Smith et al., 2004). First, FA images were created by fitting a tensor model to the raw diffusion data using FDT, and then brain-extracted using BET (Smith, 2002). All subjects’ FA data were then aligned into the FMRIB58_FA standard space using the nonlinear registration tool FNIRT (Andersson et al., 2007b, 2007a), which uses a b-spline representation of the registration warp field (Rueckert et al., 1999). Next, the mean FA image was created and thinned to create a mean FA skeleton which represents the centres of all tracts common to the group. Each subject’s aligned FA data was then projected onto this skeleton and the resulting data fed into voxelwise cross-group statistics.

Group comparisons were run using Permutation Analysis of Linear Models (PALM) (Winkler et al., 2014) with 1,000 permutations. Two contrasts were run comparing groups on the difference in FA between Day 1 and Day 5 (i.e., ([DT, Scan 5 – Scan 1] > [SOS, Scan 5 – Scan 1]), and ([SOS, Scan 5 – Scan 1] > [DT, Scan 5 – Scan 1]). Because the Double Trouble group showed a large behavioural improvement between Scan 2 and Scan 1, a second set of contrasts was run on these days (i.e., [DT, Scan 2 – Scan 1] > [SOS, Scan 2 – Scan 1]), and ([SOS, Scan 2 – Scan 1] > [DT, Scan 2 – Scan 1]). Threshold-free cluster enhancement (TFCE) was used to find significant changes in FA, and the results were thresholded at *p* = .01, Bonferroni corrected for two comparisons. The thresholded results were then thickened for presentation using *tbss_fill*, and projected onto the FMRIB58_FA_1mm brain and the mean FA skeleton of the current data for visualization. Because the contrasts only showed whether the difference between days was bigger in one group or the other, areas of significant change were used as a mask on difference FA maps to determine the direction of the change.

To assess the amount of overlap in changes to white matter microstructure between the two tasks, a conjunction analysis was conducted using Scan 5 – Scan 1 contrasts, thresholded at *p* = .01 (corrected), for each group. A final set of analyses were run to investigate whether the FA in areas that showed group differences was predicted by percent change from the beginning to end of training. First, Scan 5 – Scan 1 difference FA maps were calculated. We then conducted correlations using *randomise* (Winkler et al., 2014), with 10,000 permutations, between scores and FA values, with search space restricted to areas that had shown a significant difference between groups from Scan 1 to Scan 5. Results of the correlations were thresholded at *p* = .01.

## 3 Results

### 3.1 Cognitive training and associated learning

To assess learning trends among participants, we plotted scores achieved during cognitive training over time (Figure 2). Curve estimation was applied to each participant’s learning trend to determine whether the data fit better to a linear or logarithmic model. The “better fit” model was defined as the model whose curve estimation regression analysis returned a greater coefficient of determination (*R*^2^) value. An ANOVA comparing the two fits within the Double Trouble group confirmed that the *R*^*2*^ associated with logarithmic models (*R*^*2*^ = 0.79) was significantly higher than linear models (*R*^*2*^ = 0.62; *F*(1,38) = 31.77, *p* < .001), indicating that logarithmic models fit the participants’ learning trend data better. For the Self-Ordered Search group, there was no significant difference between the *R*^*2*^ associated with linear models (*R*^*2*^ = 0.327) and that of the logarithmic models (*R*^*2*^ = 0.333; *F*(1,38) = 0.33, *p* = .569), indicating that the logarithmic model did not fit the data better than the linear model.

**Figure 2.**
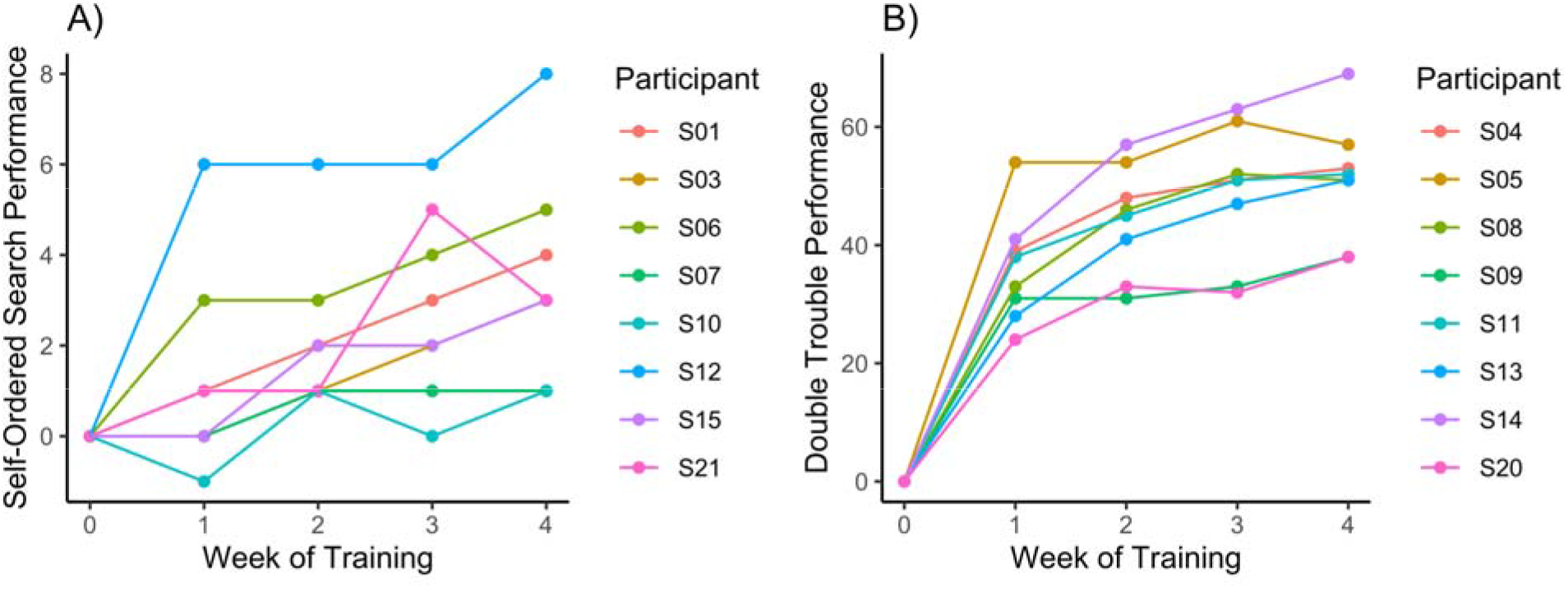
Amount of improvement across the at-home training sessions for A) Self-Ordered Search and B) Double Trouble.

We next wanted to confirm that both groups showed learning effects, and to determine whether the groups differed in the amount of learning that occurred over the cognitive training period. The results are shown in Figure 3. A linear mixed model showed a main effect of group (*F*(1,14) = 7.54, *p* = .016), a main effect of time (*F*(1,14) = 132.08, *p* < .001) and a significant group x time interaction (*F*(1,14) = 17.28, *p* < .001). Post-hoc contrasts confirmed that there was significant improvement from Week 1 to Week 5 in both the Self-Ordered Search group (*t*(14) = 5.19, *p* < .001) and the Double Trouble group (*t*(14) = 11.07, *p* < .001), and the effects of training were more pronounced for the latter group than the former.

**Figure 3.**
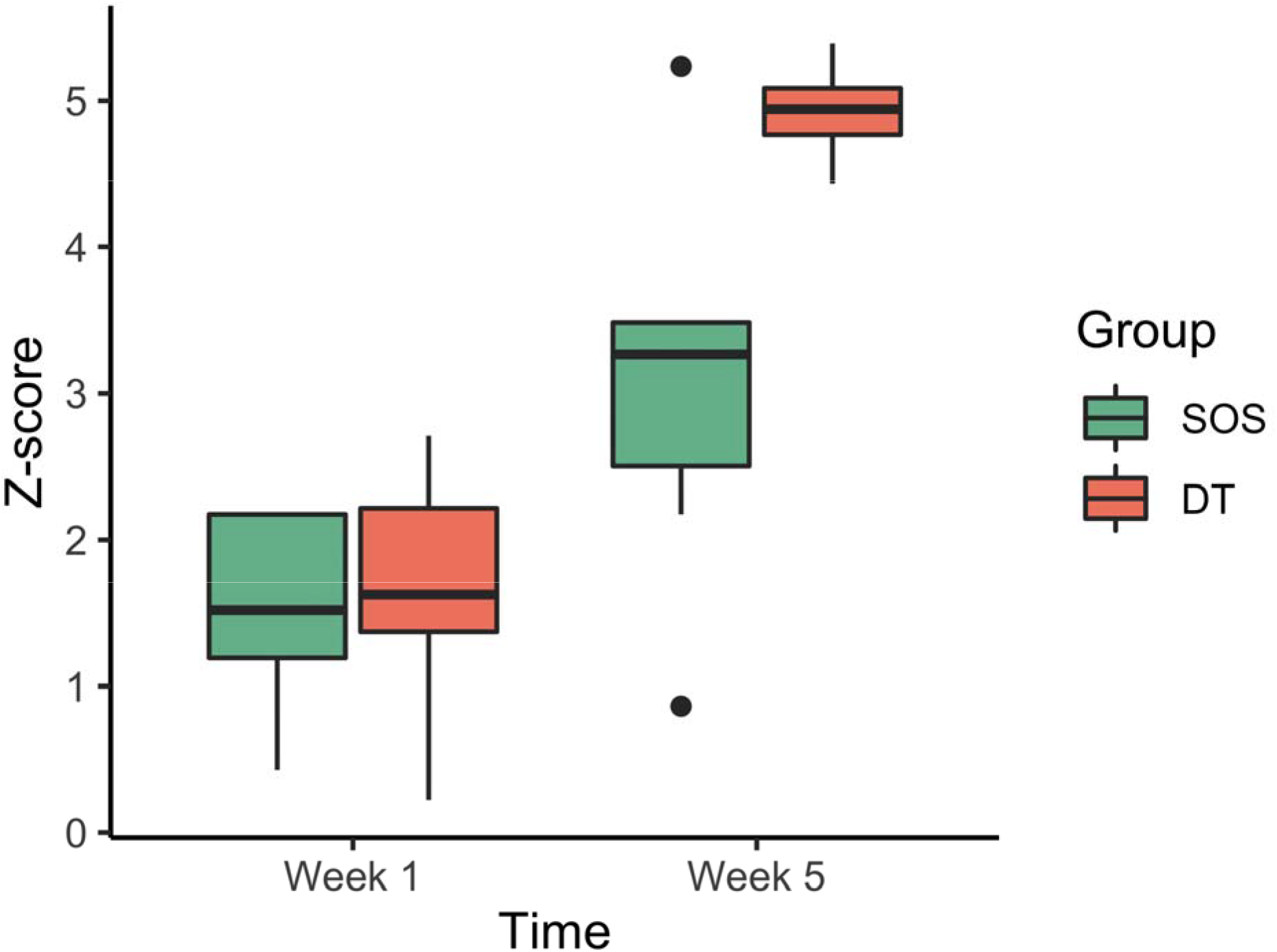
Task improvement from beginning of cognitive training to the end, for the Self-Ordered Search (SOS) and Double Trouble groups (DT). The first and third quartiles are marked by the lower and upper edges of the boxes, respectively. Lower and upper whiskers extend to the smallest and largest value, respectively, within 1.5 times the interquartile range. Outlying values beyond these ranges are plotted individually.

### 3.2 Transfer effects

We next examined whether training on Double Trouble and Self-Ordered Search led to near transfer. Grammatical Reasoning served as a measure of near transfer for those who had trained on Double Trouble and Spatial Span served as a test of near transfer for those who had trained on Self-Ordered Search. A linear mixed model indicated that there was no main effect of time (*F*(1,14) = 3.32, *p* = .090, *ns*), no main effect of training group (*F*(1,14) = 2.17, *p* = .163, *ns*), and no task x group interaction (*F*(1,14) = 0.35, *p* = .563, *ns*), indicating that no near transfer had occurred.

We then examined whether training on Double Trouble and Self-Ordered Search led to far transfer. Grammatical Reasoning served as a measure of far transfer for those who had trained on Self-Ordered Search and Spatial Span served as a test of far transfer for those who had trained on Double Trouble. A linear mixed model indicated that there was no main effect of time (*F*(1,14) = 0.01, *p* = .916, *ns*), no main effect of training group (*F*(1,14) = 0.30, *p* = .595, *ns*), and no task x group interaction (*F*(1,14) = 1.78, *p* = .203, *ns*), indicating that no far transfer had occurred.

### 3.3 DTI analysis

#### 3.3.1 Day 5 – Day 1 contrasts

To assess which regions of the white matter skeleton showed changes in FA across the entire training protocol for each group, between-groups *t*-contrasts were conducted on the difference between Scan 5 and Scan 1. Because there were a large number of significant clusters (i.e., > 300), we report here the 35 most significant tracts for each contrast, and full results are presented in the supplemental materials (Tables S1 and S2). The Double Trouble training group showed significantly greater changes than the Self-Ordered Search training group in an extensive network of regions mainly lateralized to the left hemisphere (Table 1, Figure 4). Specifically, regions of the left inferior occipitofrontal fasciculus, the uncinate, and the forceps minor showed large changes, as did the anterior thalamic radiation. A portion of the left corticospinal tract also showed large decreases in FA, which also connected to the anterior thalamic radiation. Finally, there was a large decrease in FA in the left inferior longitudinal fasciculus in the region of the temporal pole.

**Table 1.**
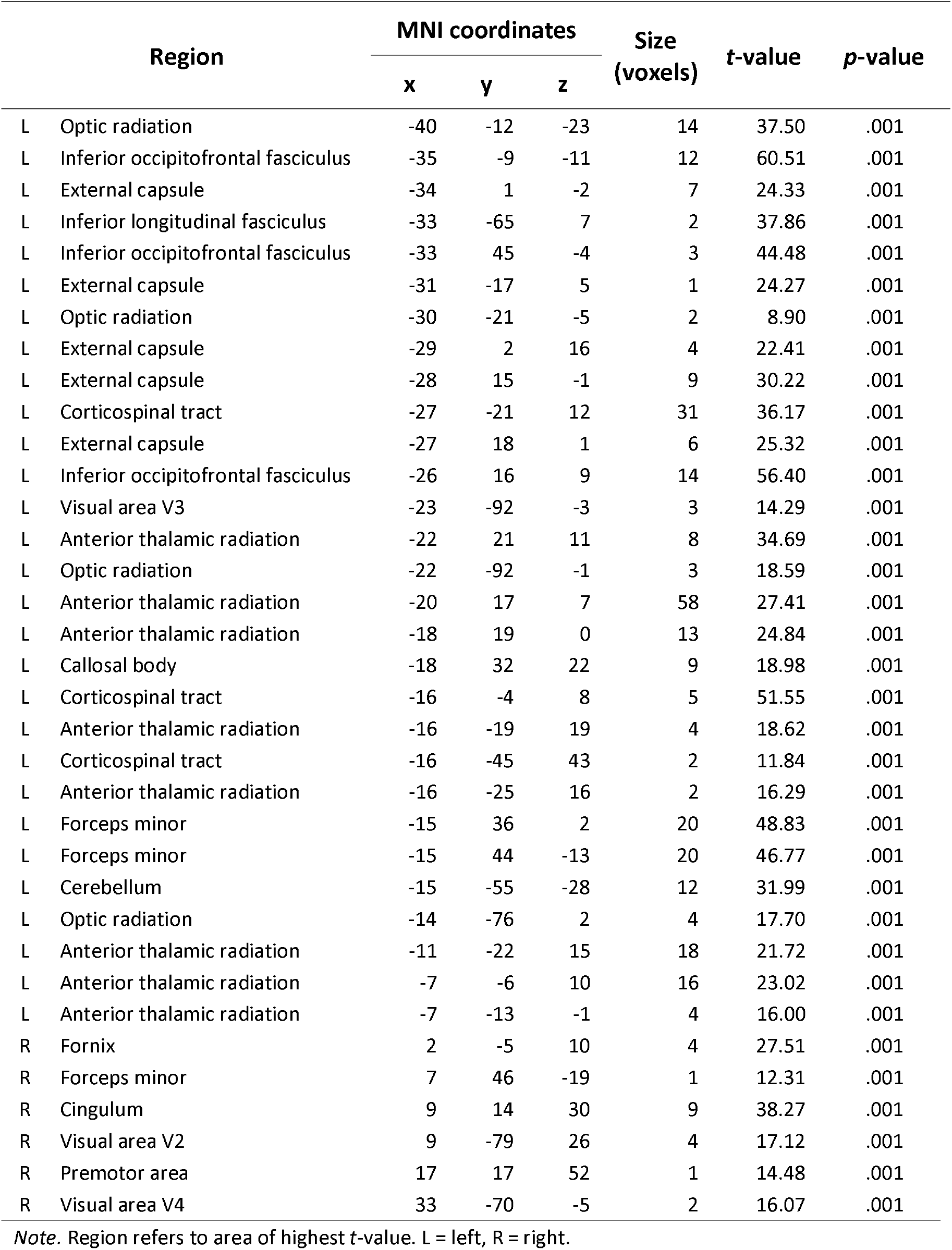
Between-group contrast of the difference between Scan 5 and Scan 1, Double Trouble > Self-Ordered Search

**Figure 4.**
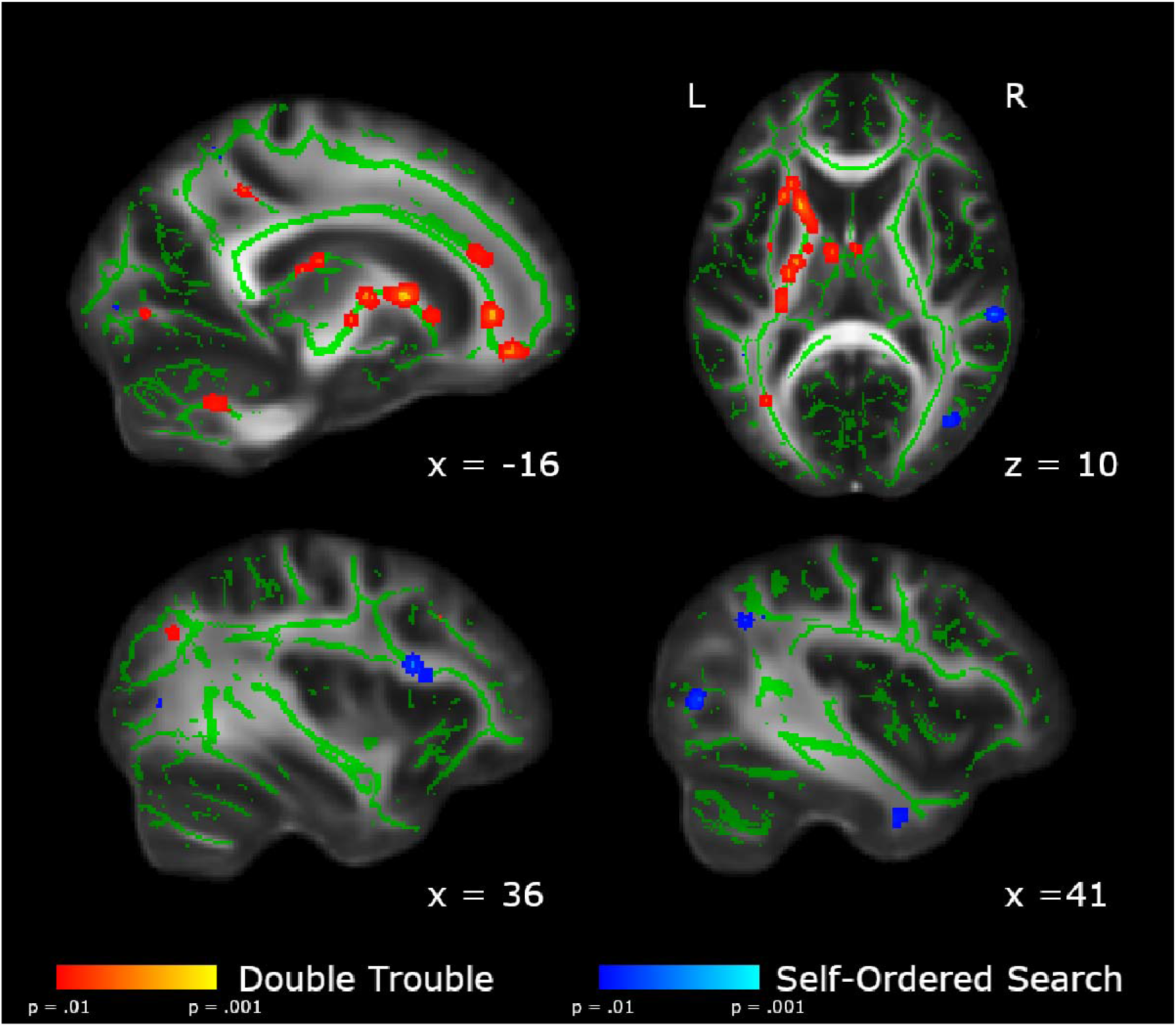
Areas showing group differences in the change in FA from Scan 1 to Scan 5. In red, we show the areas in which the changes from Scan 1 to Scan 5 are significantly larger in the Double Trouble training group than in the Self-Ordered Search training group. In blue, we show the areas in which the changes from scan 1 to scan 5 are significantly larger in the Self-Ordered Search training group than in the Double Trouble training group. Changes uniquely associated with Double Trouble were largely within the left inferior occipitofrontal and longitudinal fasciculi, while changes associated with Self-Ordered Search were largely within the right superior longitudinal fasciculus. Clusters have been thickened for visualization using *tbss_fill*, and results are overlaid on the FMRIB58_FA template and the mean skeletonized FA data of the current sample.

In contrast, the Self-Ordered Search training group showed significantly greater changes in FA than the Double Trouble training group in a right-lateralized network of regions in the dorsolateral prefrontal and parietal areas of the brain (Table 2). Specifically, the frontal section of the right superior longitudinal fasciculus showed the largest changes. There were also significant decreases in FA in a posterior section of the left superior longitudinal fasciculus within the parietal lobe, as well as the right corticospinal tract underlying the supplementary motor cortex.

**Table 2.**
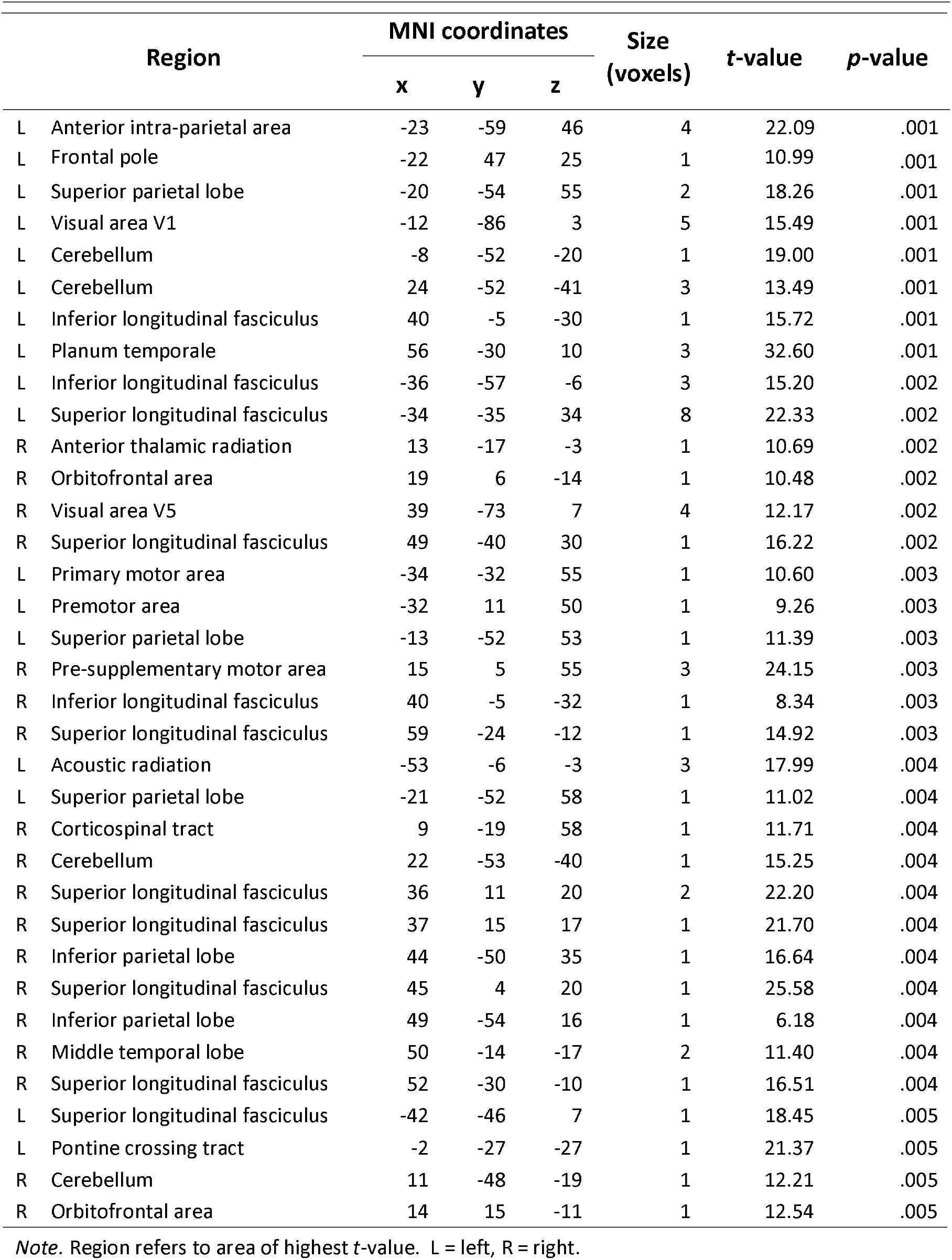
Between-group contrast of the difference between Scan 5 and Scan 1, Self-Ordered Search > Double Troble

#### 3.3.2 Day 2 – Day 1 contrasts

Because the Double Trouble training group showed a large improvement in behavioural scores within the first week, we also ran between-group contrasts on the Scan 2 – Scan 1 differences in FA (Table 3, Figure 5). The Double Trouble training group again showed left-lateralized changes, primarily in the inferior occipitofrontal fasciculus, uncinate, and forceps minor. There were also more extensive changes in the left ILF than those in Scan 5. The left forceps major also showed extensive changes in FA, extending towards, but not crossing the corpus callosum.

**Table 3.**
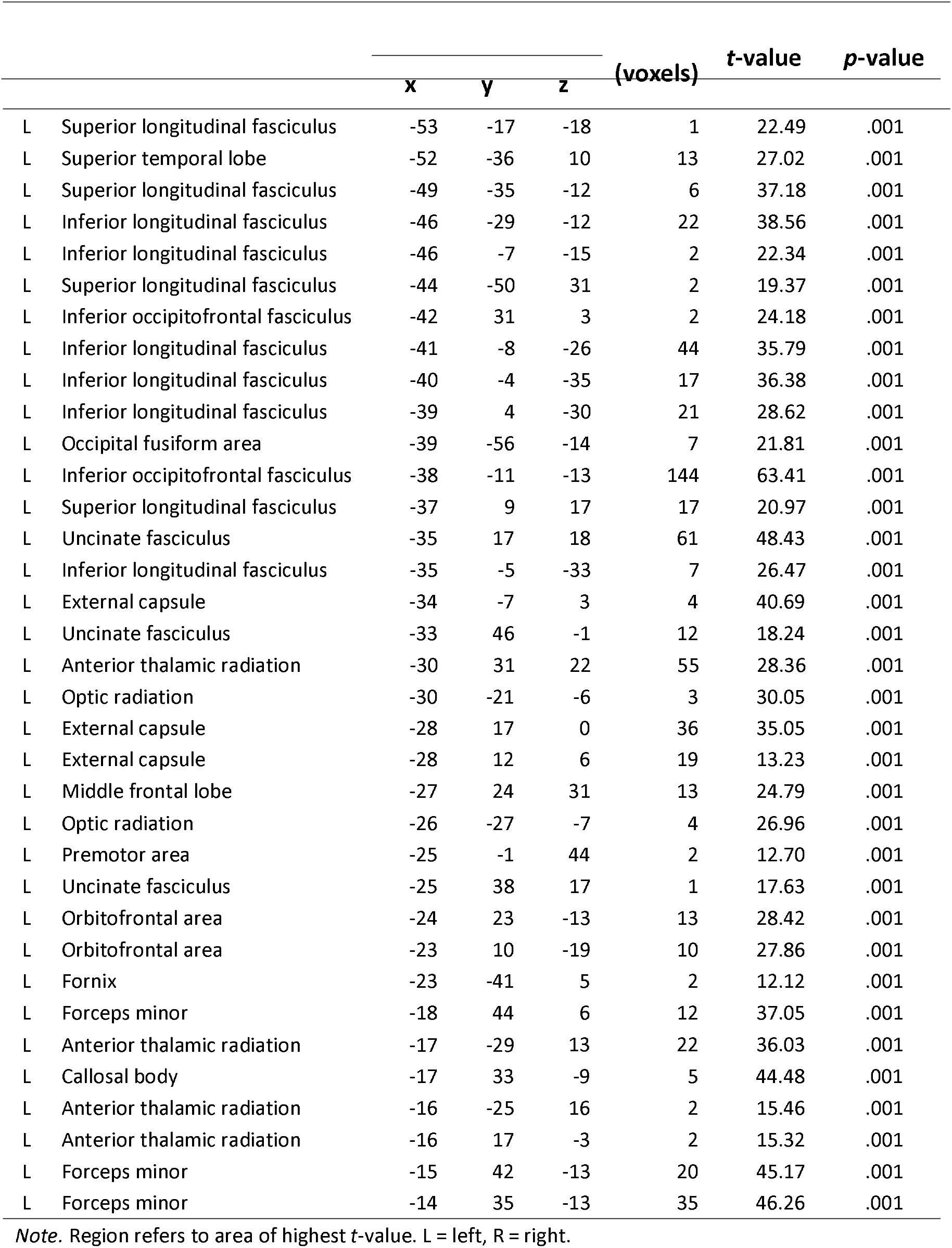
Between-group contrast of the difference between Scan 2 and Scan 1, Double Troble > Self-Ordered

**Figure 5.**
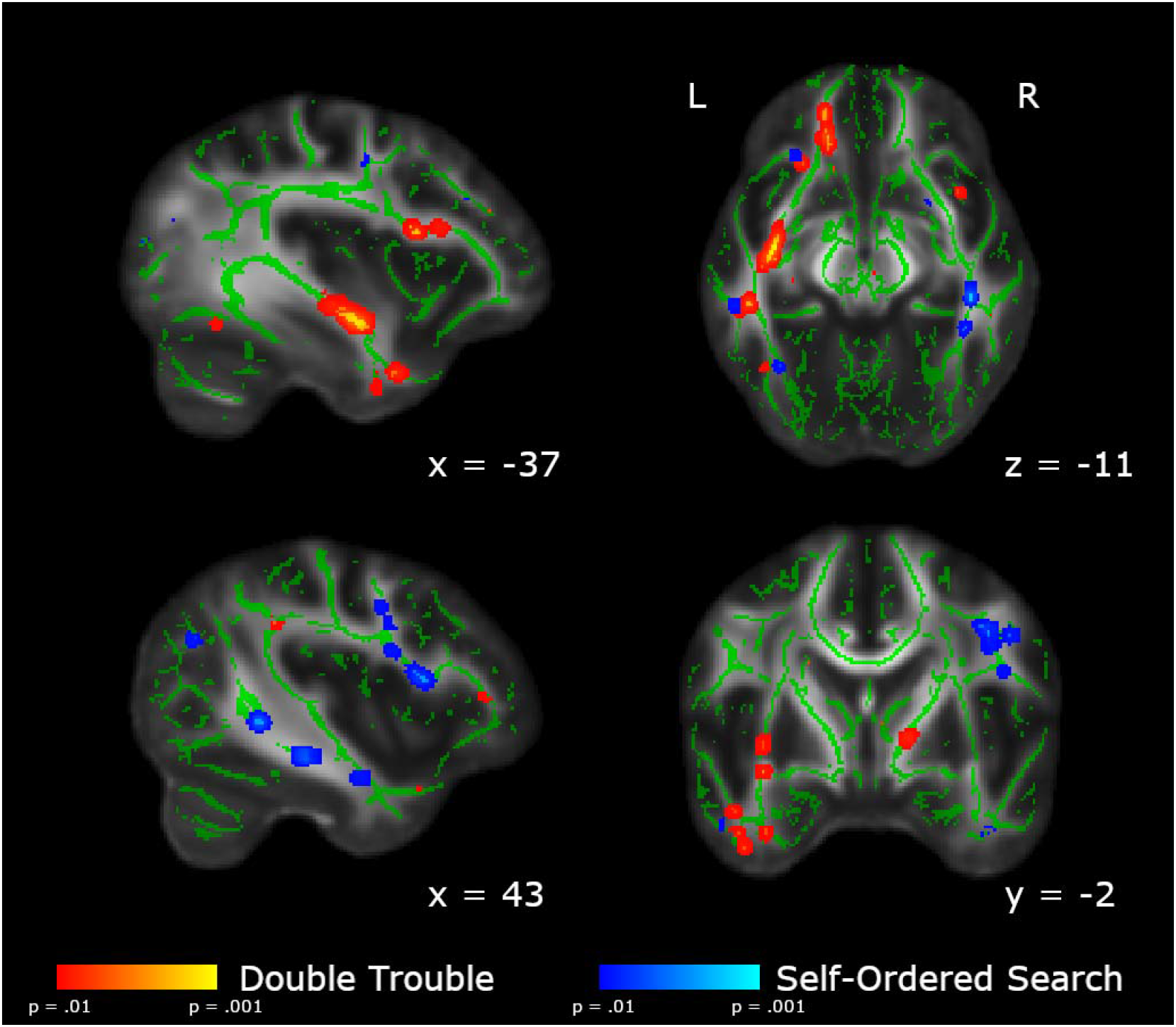
Areas showing group differences in the change in FA from Scan 1 to Scan 2. In red, we show the areas in which the changes from Scan 1 to Scan 2 are significantly larger in the Double Trouble training group than in the Self-Ordered Search training group. In blue, we show the areas in which the changes from Scan 1 to Scan 2 are significantly larger in the Self-Ordered Search training group than in the Double Trouble training group. Clusters have been thickened for visualization using *tbss_fill*, and results are overlaid on the FMRIB58_FA template and the mean skeletonized FA data of the current sample.

The Self-Ordered Search training group again showed a large cluster of changes in FA in the right dorsolateral prefrontal area of the superior longitudinal fasciculus and the posterior temporal SLF, as well as the white matter underlying the right parietal lobe (Table 4). Additionally, there was a significant cluster in the right body of the corpus callosum, however it did not cross the midline.

**Table 4.**
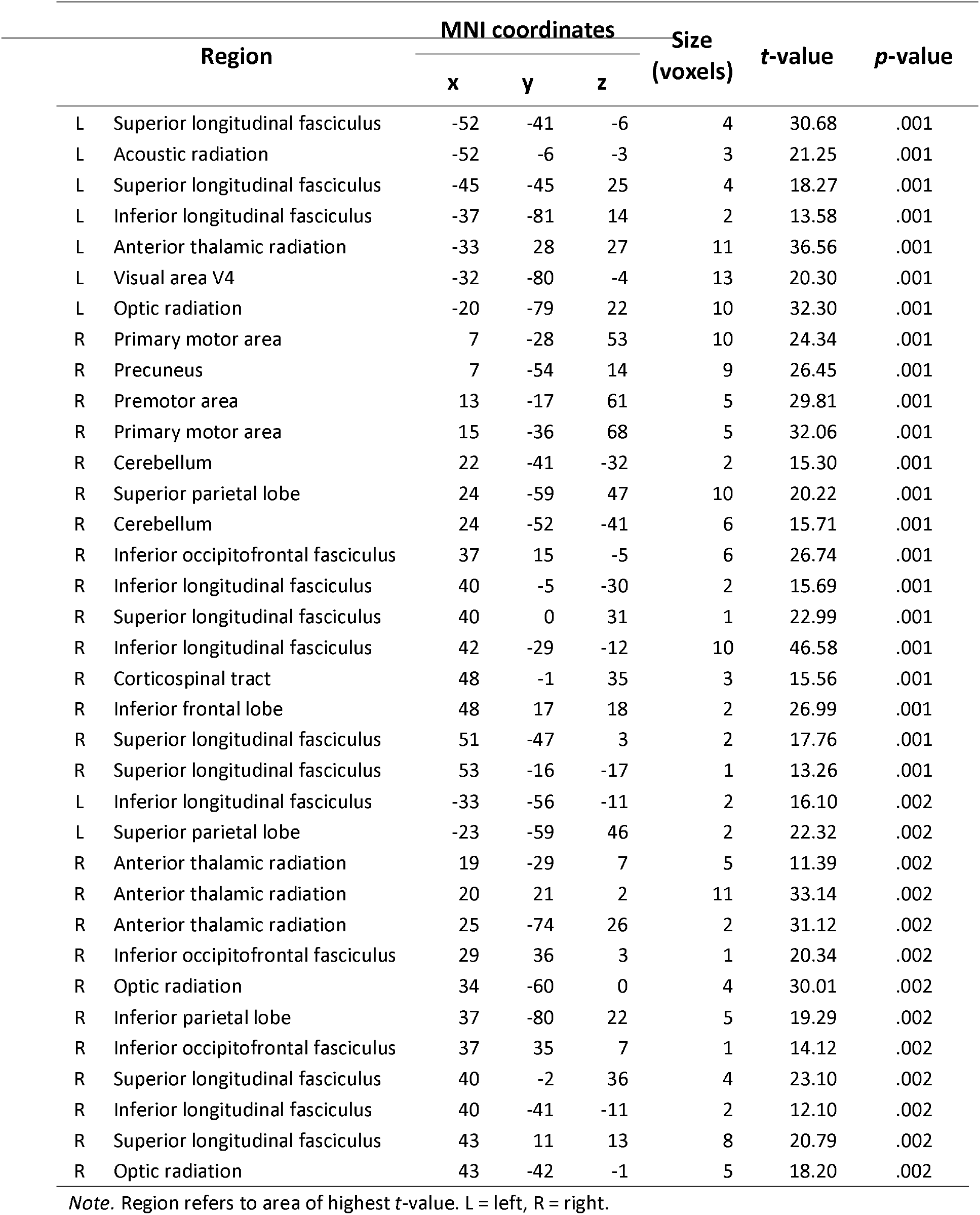
Between-group contrast of the difference between Scan 2 and Scan 1, Self-Ordered Search > Double Troble

#### 3.3.3 Conjunction analysis

To assess the degree of overlap in the changes to white matter microstructure over the course of training between the two groups, we performed a conjunction analysis. Results are shown in Figure 6 and Table 5. Very few tracts showed significant overlap between Double Trouble and Self-Ordered Search groups, including the corticospinal tract and tracts underlying the primary auditory cortex. Additionally, sections of the anterior thalamic radiation and inferior occipitofrontal fasciculus overlapped between groups, as did small regions within the superior parietal lobe and the forceps major.

**Fig 6.**
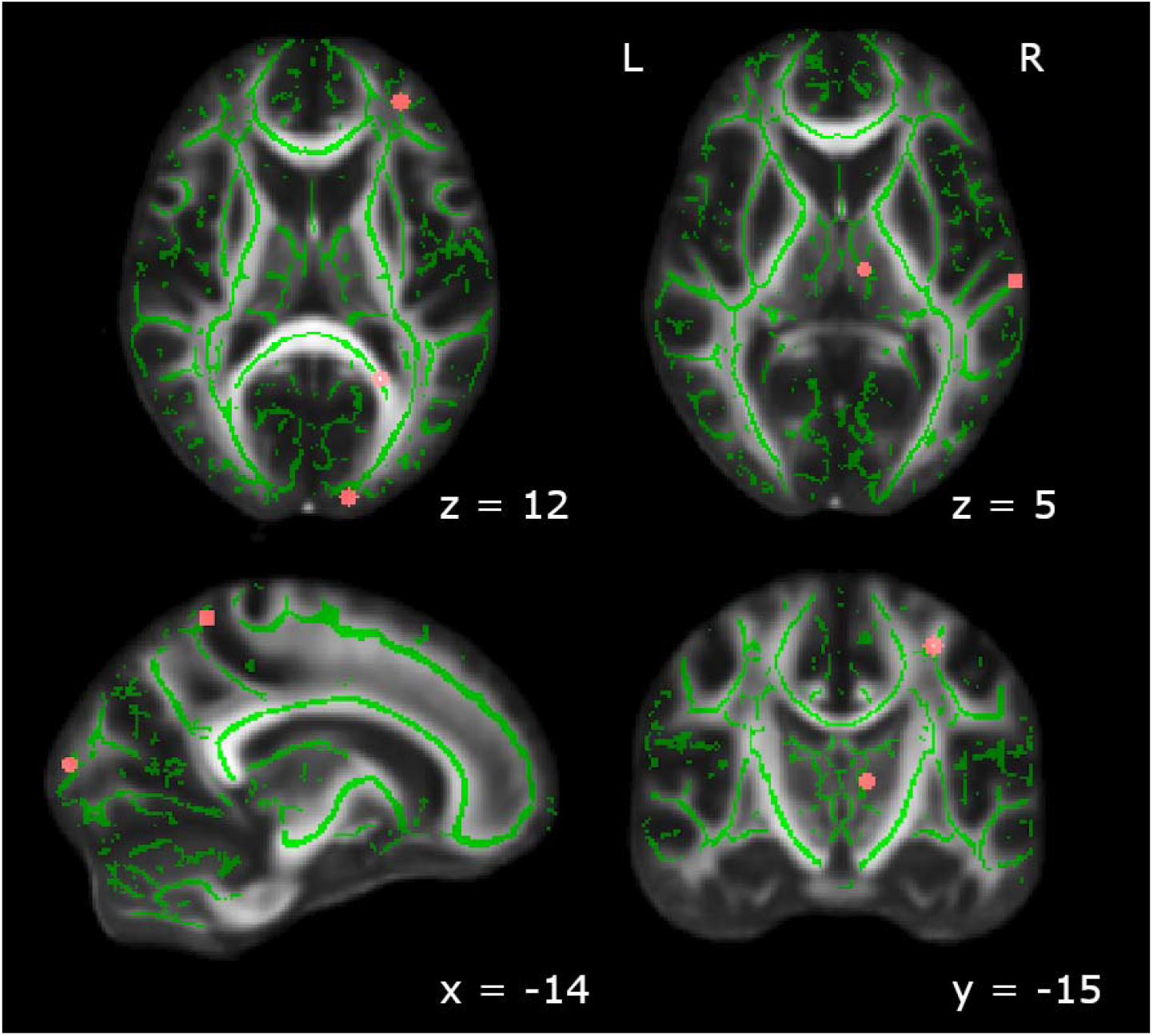
Brain regions that showed significant changes from Scan 1 to Scan 5 for both Double Trouble and Self-Ordered Search. Significant regions included the primary auditory area, the anterior thalamic radiation, the corticospinal tract, and the forceps major, as well as one region each within the inferior occipitofrontal fasciculus and the superior parietal lobe. Clusters have been thickened for visualization using *tbss_fill*, and results are overlaid on the FMRIB58_FA template and the mean skeletonized FA data of the current sample.

**Table 5.**
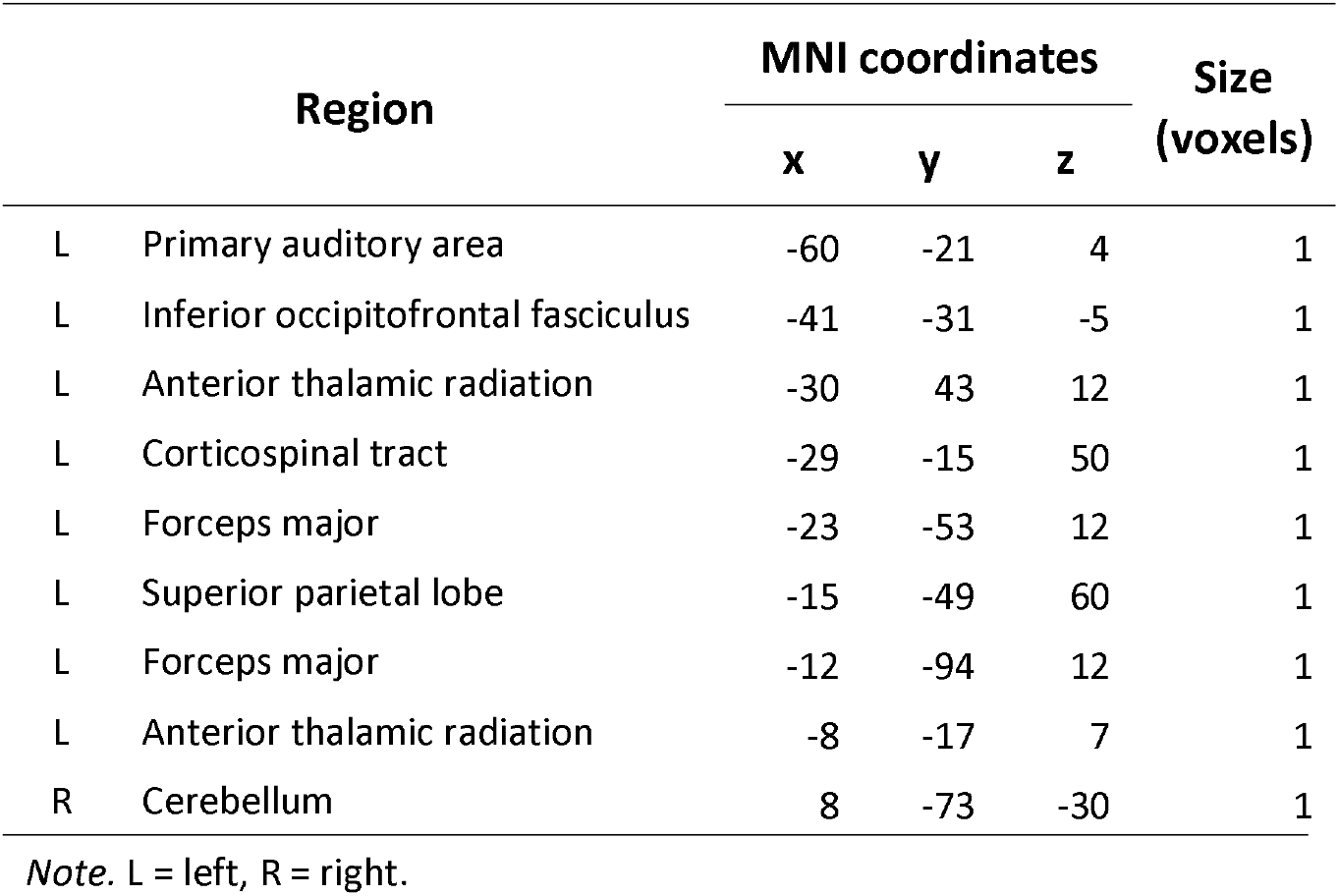
Conjunction analysis of Scan 5 – Scan 1 contrasts for Double Trouble and Self-Ordered Search training groups

| Region |  | MNI coordinates |  |  | Size<br>(voxels) |
| --- | --- | --- | --- | --- | --- |
|  |  | x | y | z |  |
| L | Primary auditory area | -60 | -21 | 4 | 1 |
| L | Inferior occipitofrontal fasciculus | -41 | -31 | -5 | 1 |
| L | Anterior thalamic radiation | -30 | 43 | 12 | 1 |
| L | Corticospinal tract | -29 | -15 | 50 | 1 |
| L | Forceps major | -23 | -53 | 12 | 1 |
| L | Superior parietal lobe | -15 | -49 | 60 | 1 |
| L | Forceps major | -12 | -94 | 12 | 1 |
| L | Anterior thalamic radiation | -8 | -17 | 7 | 1 |
| R | Cerebellum | 8 | -73 | -30 | 1 |

*Note.* L = left, R = right.

#### 3.3.4 Correlations between FA and behavioural score

To assess whether FA changes between scan 1 and scan 5 were predicted by behavioural difference scores, we conducted Pearson correlations for each group. As can be seen in Figure 7 and Table 6, several areas showed a correlation with difference scores in each group. Double Trouble performance correlated mainly with FA changes in the left hemisphere. Significant correlations existed in association fibers between several regions of the cortex including the bilateral inferior occipitofrontal fasciculus, the left superior longitudinal fasciculus, and the left uncinate fasciculus. Several commissural fibers also showed significant correlations with performance, specifically the left genu and splenium of the corpus callosum, in addition to motor pathways including white matter underlying the primary motor area.

**Figure 7.**
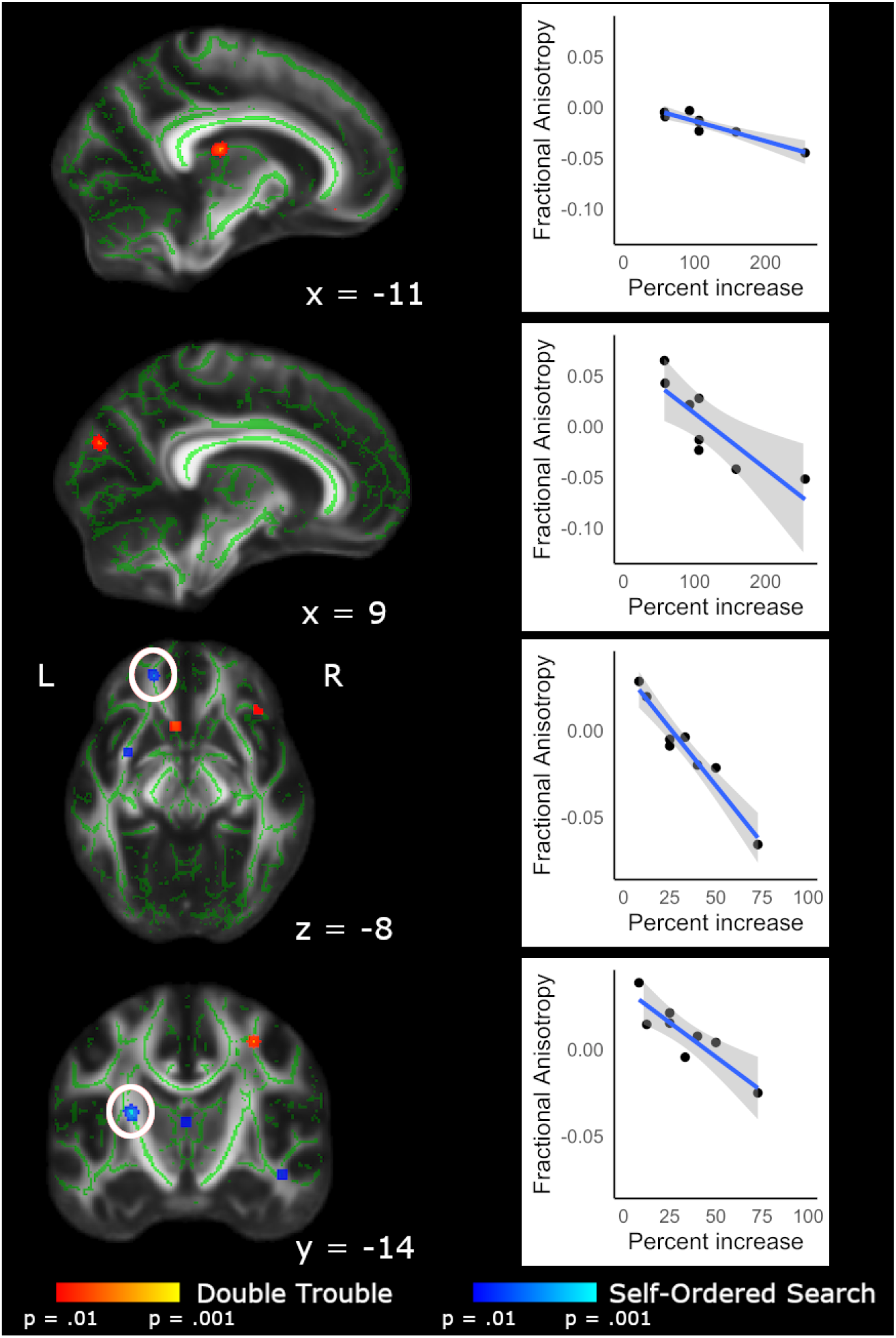
FA values from Scan 5 – Scan 1 difference maps, correlated with Scan 5 – Scan 1 difference scores. Within the brain maps, red depicts areas in which Double Trouble scores correlated with FA changes within that training group; blue depicts areas in which the Self-Ordered Search scores correlated with FA changes within that training group. Scatterplots show the relationship between difference score on the task and change in FA from Scan 1 to Scan 5. Clusters have been thickened for visualization using *tbss_fill*, and results are overlaid on the FMRIB58_FA template and the mean skeletonized FA data of the current sample.. White circles indicate the region of interest displayed in the corresponding scatterplot.

**Table 6.**
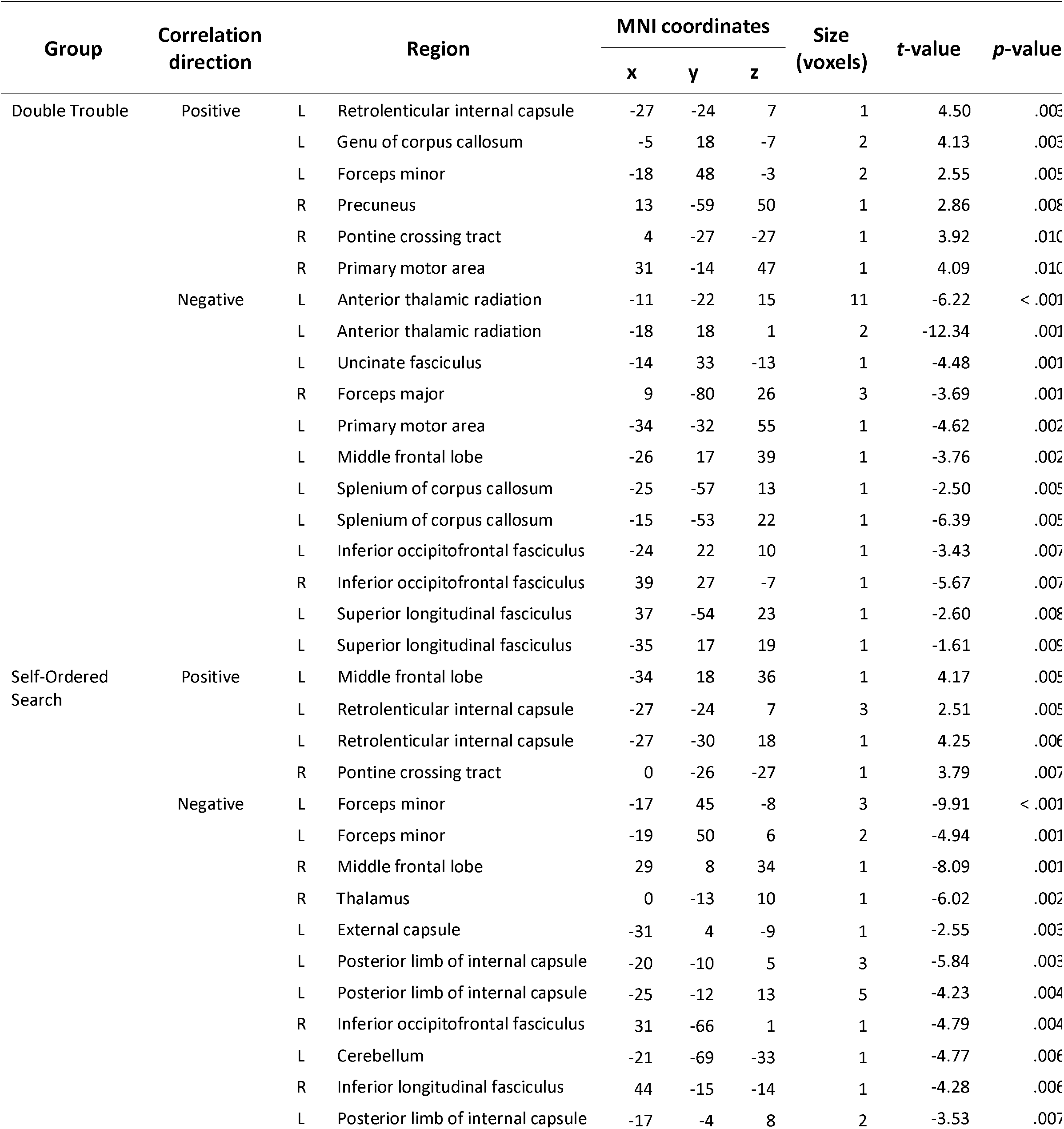
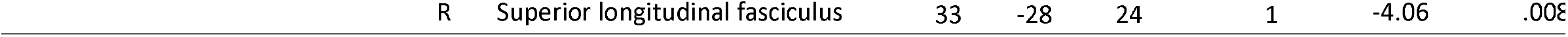
Correlations of behavioural score and FA values for Double Trouble and Self-Ordered Search training groups

Improvement on Self-Ordered Search also correlated with changes in FA in several tracts, including the bilateral superior longitudinal fasciculus. Additional correlations were seen in midbrain and frontal pathways, including the anterior thalamic radiation, thalamus, and the inferior longitudinal fasciculus, as well as in motor pathways such as the posterior limb of the internal capsule.

One voxel in the retrolenticular part of the internal capsule showed overlap between the two groups. Additionally, the two groups showed close but non-overlapping correlations in the pontine crossing tract and the forceps minor.

## 4 Discussion

In this study, we set out to investigate whether training on one of two cognitive tasks would lead to either near transfer (that is, improvements on a quantifiably similar task) or far transfer (that is, improvements on a quantifiably different task), and furthermore, if such changes exist, what the underlying neural mechanisms might be. Behaviourally, participants who trained on a spatial working memory task improved on that task over time, but did not improve on a cognitively similar test of spatial span, nor a cognitively dissimilar test of grammatical reasoning. Likewise, participants who trained on a test of inhibitory control improved on that task, but did not improve on a related test of grammatical reasoning, nor a cognitively dissimilar test of spatial span. As such, these results add to the body of work demonstrating that brain training does not ‘work’ in the sense that improvements with training in young healthy participants on cognitive tasks do not appear to generalize to other cognitive domains (Owen et al., 2010; Stojanoski et al., 2018, 2020). What then, might be a mechanistic explanation for why such training affords no generalized cognitive advantages?

To address this question, we examined changes to white matter microstructure (by measuring FA) over the course of five scanning sessions spread over the training period of four weeks. As participants trained and behaviourally improved on the primary tasks, significant changes in FA were observed in both participant groups (those that trained on Self-Ordered Search and those that trained on Double Trouble) over the course of the training period. Specifically, relative to training on Double Trouble, training on Self-Ordered Search revealed changes in integrity in the superior longitudinal fasciculus and other white matter tracts underlying frontal and parietal areas of the brain, particularly in the right hemisphere. This task has been shown to be highly sensitive to neurosurgical excisions of the frontal lobe (Owen et al., 1990) and specifically activates the mid-dorsolateral prefrontal cortex and the posterior parietal cortex in healthy participants (Owen et al., 1996). Moreover, the role of the right mid-dorsolateral prefrontal cortex has been shown to be in the involvement of task-specific strategies that lead to improvements in performance through the adoption of a repetitive searching pattern of behaviour (Owen et al., 1990, 1996). It is perhaps then not surprising that repetitive training on this task leads to white matter changes in a network of tracts that connect and support the functioning of these two regions.

In contrast, training on Double Trouble, a word-based test of inhibitory control, led to white matter changes that were predominantly in the left hemisphere and included the inferior longitudinal fasciculus and the longitudinal occipitofrontal fasciculus. A substantial literature exists detailing the role of ventral and orbitofrontal regions in tests of inhibitory control (Bryden & Roesch, 2015; Elliott & Deakin, 2005; He et al., 2019; Horn et al., 2003; Stuss et al., 2001; Szatkowska et al., 2007). Patients with damage to the orbitofrontal cortex also exhibit failures of inhibitory control, including OCD (Abe et al., 2015; Maia et al., 2008), obsessive gambling behaviour (Cavedini et al., 2002), alcoholism (Medina et al., 2008; Volkow et al., 1993), and sexual disinhibition (Gorman & Cummings, 1992; Miller et al., 1986). There is also a substantial literature in non-human primates detailing the relationship between orbitofrontal lesions and failures of inhibitory control (McEnaney & Butter, 1969; Oikonomidis et al., 2017; Wallis et al., 2001). Again, the fact that we observed white matter changes in tracts that support connectivity to this region while participants were training and improving on a test of inhibition is therefore perhaps not surprising.

Remarkably, there was almost no overlap between the white matter changes that were observed in the tracts that support improvements on Self-Ordered Search and those that support improvements on Double Trouble. In fact, a formal conjunction analysis revealed no regions with more than a single voxel in common to both training regimens, and even then, those changes were primarily in auditory, thalamic, and visual regions. Moreover, when we examined those regions in which white matter changes correlated with performance improvements, there was again very little overlap between the two training regimens.

Of course, in the description above of white matter changes that were associated with training on each of the two tasks, we focused mainly on regions that are known to be functionally involved in those tasks. In both cases, many other tracts showed changes (in some cases, over 200 areas, see supplementary materials Tables S1-S4). Nevertheless, the important point is that there was virtually no overlap between these two sets of regions as indexed by the conjunction analysis, which revealed almost no common areas of change.

Therefore, on the basis of these findings, we propose that training on one cognitive task for four weeks does not lead to improvements on a cognitively dissimilar task because the underlying white matter tracts that support communication between regions involved in those tasks are almost completely non-overlapping. Put simply, improvements in a test of spatial working memory with training are underpinned by changes in a task-specific network of brain regions that are not involved in supporting other tests like those of inhibitory control, and vice versa. The fact that we did not see any improvements in the tests of “near transfer” (i.e., Spatial Span for the Self-Ordered Search task, and Grammatical Reasoning for the Double Trouble task) suggests further that even tasks that have substantial and quantifiable similarities in the regions that they recruit (Hampshire et al., 2012) do not benefit from the white matter changes induced by training on a different task. If correct, this means that the white matter changes associated with training on any given cognitive task are so specific to that task that they do not lead to improvements on a conceptually similar task.

While these findings explain why we did not find any near or far transfer in this study, they do not explain why some other studies have reported such effects (Au et al., 2015; Caeyenberghs et al., 2016; Jaeggi et al., 2008). The variability in results seen across studies likely relates to inconsistent and often vague definitions of what constitutes ‘transfer’. The terms ‘near’ and ‘far’ transfer are often used to refer to improvements in closely related and unrelated cognitive tasks, respectively, yet how ‘related’ one task actually is to another is rarely quantified. Tasks are often selected based on their inferred cognitive properties, rather than on an empirical measure of similarity, and without a consistent definition of transfer, and quantifiable measures of similarity between tasks, it is very difficult to make comparisons across studies, and assess the reliability of any observed training-related benefits. Of course, an argument could be made that with a longer period of training, or with an increased number of participants, we may have found some evidence of near, or even far, transfer in this study. However, given the marked differences between the white matter changes associated with training on these two cognitively different tasks over 4 weeks, it seems very unlikely that increasing the length of training or the number of participants would fundamentally alter that emerging pattern.

A final point of note is that not all of the changes in FA that correlated with training were in the same direction (see Figure 7). FA quantifies the degree of cohesiveness of white matter tracts, based on the degree of diffusion of water molecules. While higher FA was originally thought to represent ‘better’ white matter integrity, numerous studies have found decreases with experience or training (Nichols & Joanisse, 2016). Although it is difficult to interpret the direction of the correlations in the present study, we can nevertheless confirm that these regions showed *change* in white matter microstructure with training.

In conclusion, in this study we showed that improvements through training on a cognitive task for four weeks do not transfer to other cognitive tasks, even those that are quantifiably similar. We suggest that this lack of near or far transfer occurs because changes in white matter tracts associated with training on each task are almost entirely non-overlapping, and therefore afford no advantages for untrained tasks.

## Acknowledgments

We are grateful to Anderson M. Winkler for help with the DTI analysis using PALM.

