## Supplemental materials for "Longitudinal white matter changes associated with cognitive training"

| **Table S1. Between-group contrast of the difference between Scan 5 and Scan 1, Double Trouble > Self-Ordered Search, full list of significant regions** | | | | | | | |
| --- | --- | --- | --- | --- | --- | --- | --- |
| **Region** | | **MNI coordinates** | | | **Size (voxels)** | ***t*-value** | ***p*-value** |
|  |  | **x** | **y** | **z** |  |  |  |
| L | Optic radiation | -40 | -12 | -23 | 14 | 37.50 | .001 |
| L | Inferior occipitofrontal fasciculus | -35 | -9 | -11 | 12 | 60.51 | .001 |
| L | External capsule | -34 | 1 | -2 | 7 | 24.33 | .001 |
| L | Inferior longitudinal fasciculus | -33 | -65 | 7 | 2 | 37.86 | .001 |
| L | Inferior occipitofrontal fasciculus | -33 | 45 | -4 | 3 | 44.48 | .001 |
| L | External capsule | -31 | -17 | 5 | 1 | 24.27 | .001 |
| L | Optic radiation | -30 | -21 | -5 | 2 | 8.90 | .001 |
| L | External capsule | -29 | 2 | 16 | 4 | 22.41 | .001 |
| L | External capsule | -28 | 15 | -1 | 9 | 30.22 | .001 |
| L | Corticospinal tract | -27 | -21 | 12 | 31 | 36.17 | .001 |
| L | External capsule | -27 | 18 | 1 | 6 | 25.32 | .001 |
| L | Inferior occipitofrontal fasciculus | -26 | 16 | 9 | 14 | 56.40 | .001 |
| L | Visual area V3 | -23 | -92 | -3 | 3 | 14.29 | .001 |
| L | Anterior thalamic radiation | -22 | 21 | 11 | 8 | 34.69 | .001 |
| L | Optic radiation | -22 | -92 | -1 | 3 | 18.59 | .001 |
| L | Anterior thalamic radiation | -20 | 17 | 7 | 58 | 27.41 | .001 |
| L | Anterior thalamic radiation | -18 | 19 | 0 | 13 | 24.84 | .001 |
| L | Callosal body | -18 | 32 | 22 | 9 | 18.98 | .001 |
| L | Corticospinal tract | -16 | -4 | 8 | 5 | 51.55 | .001 |
| L | Anterior thalamic radiation | -16 | -19 | 19 | 4 | 18.62 | .001 |
| L | Corticospinal tract | -16 | -45 | 43 | 2 | 11.84 | .001 |
| L | Anterior thalamic radiation | -16 | -25 | 16 | 2 | 16.29 | .001 |
| L | Forceps minor | -15 | 36 | 2 | 20 | 48.83 | .001 |
| L | Forceps minor | -15 | 44 | -13 | 20 | 46.77 | .001 |
| L | Cerebellum | -15 | -55 | -28 | 12 | 31.99 | .001 |
| L | Optic radiation | -14 | -76 | 2 | 4 | 17.70 | .001 |
| L | Anterior thalamic radiation | -11 | -22 | 15 | 18 | 21.72 | .001 |
| L | Anterior thalamic radiation | -7 | -6 | 10 | 16 | 23.02 | .001 |
| L | Anterior thalamic radiation | -7 | -13 | -1 | 4 | 16.00 | .001 |
| R | Fornix | 2 | -5 | 10 | 4 | 27.51 | .001 |
| R | Forceps minor | 7 | 46 | -19 | 1 | 12.31 | .001 |
| R | Cingulum | 9 | 14 | 30 | 9 | 38.27 | .001 |
| R | Visual area V2 | 9 | -79 | 26 | 4 | 17.12 | .001 |
| R | Premotor area | 17 | 17 | 52 | 1 | 14.48 | .001 |
| R | Visual area V4 | 33 | -70 | -5 | 2 | 16.07 | .001 |
| R | Inferior occipitofrontal fasciculus | 34 | 49 | -4 | 1 | 14.21 | .001 |
| L | Corticospinal tract | -47 | -2 | 33 | 1 | 12.99 | .002 |
| L | Inferior frontal gyrus | -42 | 14 | 23 | 2 | 13.47 | .002 |
| L | Uncinate fasciculus | -38 | 10 | -32 | 3 | 17.28 | .002 |
| L | Uncinate fasciculus | -36 | 37 | -5 | 4 | 20.28 | .002 |
| L | External capsule | -32 | -3 | 9 | 2 | 21.43 | .002 |
| L | Optic radiation | -32 | -78 | 1 | 1 | 16.99 | .002 |
| L | Callosal body | -29 | -48 | 22 | 2 | 15.62 | .002 |
| L | Corticospinal tract | -24 | -13 | 13 | 21 | 55.03 | .002 |
| L | Corticospinal tract | -21 | -10 | 10 | 11 | 42.93 | .002 |
| L | Superior parietal lobe | -19 | -43 | 40 | 5 | 44.54 | .002 |
| L | Anterior corona radiata | -17 | 25 | -12 | 1 | 8.50 | .002 |
| L | Callosal body | -13 | 21 | -9 | 1 | 8.70 | .002 |
| L | Fornix | -5 | 2 | -7 | 2 | 12.80 | .002 |
| R | Anterior thalamic radiation | 4 | -5 | 7 | 1 | 15.45 | .002 |
| R | Anterior thalamic radiation | 5 | -3 | 6 | 4 | 18.20 | .002 |
| R | Anterior thalamic radiation | 10 | -19 | 8 | 1 | 16.77 | .002 |
| R | Frontal pole | 37 | 36 | 18 | 1 | 12.21 | .002 |
| L | Planum temporale | -55 | -21 | 2 | 3 | 22.21 | .003 |
| L | Inferior longitudinal fasciculus | -50 | -6 | -23 | 1 | 15.99 | .003 |
| L | Inferior longitudinal fasciculus | -43 | 0 | -23 | 2 | 15.78 | .003 |
| L | Inferior occipitofrontal fasciculus | -37 | -13 | -12 | 3 | 29.94 | .003 |
| L | Inferior occipitofrontal fasciculus | -35 | -3 | -5 | 3 | 15.80 | .003 |
| L | Callosal body | -33 | -64 | 9 | 2 | 24.65 | .003 |
| L | Inferior occipitofrontal fasciculus | -25 | 21 | 3 | 5 | 31.29 | .003 |
| L | Inferior occipitofrontal fasciculus | -24 | 31 | 2 | 10 | 42.98 | .003 |
| L | Inferior occipitofrontal fasciculus | -24 | 32 | -2 | 1 | 24.90 | .003 |
| L | Uncinate fasciculus | -19 | 29 | -9 | 10 | 33.15 | .003 |
| L | Inferior occipitofrontal fasciculus | -19 | 54 | -11 | 1 | 12.52 | .003 |
| L | Anterior thalamic radiation | -9 | -20 | 7 | 2 | 8.86 | .003 |
| R | Primary motor area | 7 | -33 | 66 | 1 | 23.79 | .003 |
| R | Fornix | 9 | -29 | 16 | 2 | 15.37 | .003 |
| R | Corticospinal tract | 13 | -24 | -17 | 6 | 36.58 | .003 |
| R | Middle frontal gyrus | 36 | 20 | 36 | 1 | 13.57 | .003 |
| R | Inferior parietal lobe | 37 | -69 | 31 | 3 | 20.17 | .003 |
| L | Inferior longitudinal fasciculus | -44 | -9 | -19 | 1 | 18.68 | .004 |
| L | Temporal occipital fusiform area | -40 | -57 | -14 | 2 | 16.46 | .004 |
| L | Superior longitudinal fasciculus | -40 | -12 | 28 | 2 | 38.04 | .004 |
| L | Inferior occipitofrontal fasciculus | -37 | -11 | -13 | 1 | 25.15 | .004 |
| L | Optic radiation | -33 | -63 | 13 | 3 | 21.08 | .004 |
| L | Inferior occipitofrontal fasciculus | -33 | -5 | -9 | 2 | 27.53 | .004 |
| L | Middle frontal lobe | -30 | 27 | 36 | 1 | 7.61 | .004 |
| L | Anterior thalamic radiation | -22 | 33 | 26 | 1 | 33.54 | .004 |
| L | Anterior thalamic radiation | -21 | 34 | 24 | 2 | 36.17 | .004 |
| L | Forceps minor | -20 | 42 | 20 | 5 | 37.25 | .004 |
| L | Forceps minor | -18 | 48 | -3 | 2 | 23.31 | .004 |
| L | Forceps minor | -18 | 50 | 6 | 2 | 14.46 | .004 |
| L | Optic radiation | -17 | -61 | 1 | 1 | 12.60 | .004 |
| L | Corticospinal tract | -16 | -2 | 7 | 1 | 47.83 | .004 |
| L | Anterior thalamic radiation | -14 | -2 | 4 | 2 | 31.21 | .004 |
| L | Forceps minor | -13 | 52 | -10 | 1 | 22.51 | .004 |
| L | Anterior thalamic radiation | -11 | -4 | -4 | 2 | 31.42 | .004 |
| L | Anterior thalamic radiation | -10 | -20 | 11 | 1 | 15.79 | .004 |
| L | Fornix | -7 | -24 | 16 | 2 | 17.28 | .004 |
| L | Brainstem | -5 | -30 | -23 | 2 | 35.82 | .004 |
| R | Anterior thalamic radiation | 4 | -12 | -10 | 1 | 14.88 | .004 |
| R | Brainstem | 6 | -16 | -24 | 1 | 21.24 | .004 |
| R | Fornix | 9 | -26 | 4 | 1 | 10.55 | .004 |
| R | Superior parietal lobe | 12 | -59 | 50 | 1 | 8.88 | .004 |
| R | Cerebellum | 22 | -56 | -32 | 1 | 27.10 | .004 |
| R | Cerebellum | 22 | -58 | -32 | 1 | 28.53 | .004 |
| R | Orbitofrontal area | 25 | 14 | -19 | 1 | 16.99 | .004 |
| R | Middle frontal lobe | 29 | 8 | 34 | 1 | 12.00 | .004 |
| R | Visual area V4 | 33 | -68 | -8 | 1 | 12.39 | .004 |
| R | Superior longitudinal fasciculus | 35 | -47 | 25 | 1 | 8.50 | .004 |
| R | Superior longitudinal fasciculus | 37 | -54 | 23 | 2 | 14.14 | .004 |
| R | Temporal fusiform area | 38 | -36 | -18 | 3 | 15.23 | .004 |
| R | Frontal pole | 38 | 36 | 20 | 1 | 10.72 | .004 |
| R | Inferior longitudinal fasciculus | 45 | -15 | -14 | 2 | 24.52 | .004 |
| L | Inferior longitudinal fasciculus | -44 | -11 | -16 | 1 | 17.90 | .005 |
| L | Superior longitudinal fasciculus | -42 | 10 | 13 | 1 | 18.41 | .005 |
| L | External capsule | -32 | 1 | 7 | 1 | 11.19 | .005 |
| L | Hippocampus | -32 | -36 | -12 | 1 | 14.22 | .005 |
| L | Inferior occipitofrontal fasciculus | -31 | 3 | -10 | 1 | 12.56 | .005 |
| L | Hippocampus | -30 | -32 | -15 | 1 | 13.11 | .005 |
| L | External capsule | -29 | 4 | 12 | 2 | 16.17 | .005 |
| L | Inferior occipitofrontal fasciculus | -29 | 5 | -10 | 1 | 14.69 | .005 |
| L | Optic radiation | -26 | -29 | -3 | 1 | 10.93 | .005 |
| L | Optic radiation | -26 | -28 | -7 | 1 | 12.66 | .005 |
| L | Inferior occipitofrontal fasciculus | -24 | 22 | 6 | 2 | 35.09 | .005 |
| L | Anterior thalamic radiation | -23 | 36 | 23 | 1 | 22.71 | .005 |
| L | Inferior occipitofrontal fasciculus | -20 | 51 | -9 | 1 | 8.97 | .005 |
| L | Forceps minor | -20 | 33 | 26 | 1 | 26.06 | .005 |
| L | Forceps minor | -19 | 37 | 20 | 6 | 33.71 | .005 |
| L | Forceps minor | -19 | 36 | 25 | 1 | 29.12 | .005 |
| L | Anterior thalamic radiation | -17 | -29 | 13 | 1 | 18.72 | .005 |
| L | Cingulum | -17 | 32 | 26 | 1 | 23.22 | .005 |
| L | Corticospinal tract | -16 | -9 | -1 | 2 | 31.35 | .005 |
| L | Superior parietal lobe | -15 | -42 | 41 | 1 | 14.49 | .005 |
| R | Anterior thalamic radiation | 2 | -13 | -7 | 1 | 10.23 | .005 |
| R | Anterior thalamic radiation | 10 | -20 | 0 | 1 | 14.56 | .005 |
| R | Precuneus | 13 | -57 | 51 | 1 | 11.70 | .005 |
| R | Forceps major | 15 | -83 | 25 | 1 | 9.64 | .005 |
| R | Forceps minor | 16 | 55 | 12 | 1 | 16.52 | .005 |
| R | Cingulum | 17 | -55 | 36 | 2 | 21.37 | .005 |
| R | Premotor area | 17 | 15 | 52 | 1 | 11.50 | .005 |
| R | Callosal body | 19 | 12 | 30 | 1 | 5.03 | .005 |
| R | Cerebellum | 32 | -52 | -38 | 1 | 12.30 | .005 |
| R | Anterior thalamic radiation | 32 | 34 | 16 | 1 | 16.61 | .005 |
| R | Anterior thalamic radiation | 32 | 32 | 17 | 1 | 17.63 | .005 |
| R | Middle frontal lobe | 34 | 34 | 29 | 1 | 6.60 | .005 |
| R | Superior longitudinal fasciculus | 55 | 6 | 13 | 1 | 7.77 | .005 |
| R | Planum temporale | 60 | -23 | 9 | 1 | 7.18 | .005 |
| *Note.* Region refers to area of highest *t*-value. L = left, R = right. | | | | | | | |

| **Table S2. Between-group contrast of the difference between Scan 5 and Scan 1, Self-Ordered Search > Double Trouble, full list of significant regions** | | | | | | | |
| --- | --- | --- | --- | --- | --- | --- | --- |
| **Region** | | **MNI coordinates** | | | **Size (voxels)** | ***t*-value** | ***p*-value** |
|  |  | **x** | **y** | **z** |  |  |  |
| L | Anterior intra-parietal area | -23 | -59 | 46 | 4 | 22.09 | .001 |
| L | Frontal pole | -22 | 47 | 25 | 1 | 10.99 | .001 |
| L | Superior parietal lobe | -20 | -54 | 55 | 2 | 18.26 | .001 |
| L | Visual area V1 | -12 | -86 | 3 | 5 | 15.49 | .001 |
| L | Cerebellum | -8 | -52 | -20 | 1 | 19.00 | .001 |
| L | Cerebellum | 24 | -52 | -41 | 3 | 13.49 | .001 |
| L | Inferior longitudinal fasciculus | 40 | -5 | -30 | 1 | 15.72 | .001 |
| L | Planum temporale | 56 | -30 | 10 | 3 | 32.60 | .001 |
| L | Inferior longitudinal fasciculus | -36 | -57 | -6 | 3 | 15.20 | .002 |
| L | Superior longitudinal fasciculus | -34 | -35 | 34 | 8 | 22.33 | .002 |
| R | Anterior thalamic radiation | 13 | -17 | -3 | 1 | 10.69 | .002 |
| R | Orbitofrontal area | 19 | 6 | -14 | 1 | 10.48 | .002 |
| R | Visual area V5 | 39 | -73 | 7 | 4 | 12.17 | .002 |
| R | Superior longitudinal fasciculus | 49 | -40 | 30 | 1 | 16.22 | .002 |
| L | Primary motor area | -34 | -32 | 55 | 1 | 10.60 | .003 |
| L | Premotor area | -32 | 11 | 50 | 1 | 9.26 | .003 |
| L | Superior parietal lobe | -13 | -52 | 53 | 1 | 11.39 | .003 |
| R | Pre-supplementary motor area | 15 | 5 | 55 | 3 | 24.15 | .003 |
| R | Inferior longitudinal fasciculus | 40 | -5 | -32 | 1 | 8.34 | .003 |
| R | Superior longitudinal fasciculus | 59 | -24 | -12 | 1 | 14.92 | .003 |
| L | Acoustic radiation | -53 | -6 | -3 | 3 | 17.99 | .004 |
| L | Superior parietal lobe | -21 | -52 | 58 | 1 | 11.02 | .004 |
| R | Corticospinal tract | 9 | -19 | 58 | 1 | 11.71 | .004 |
| R | Cerebellum | 22 | -53 | -40 | 1 | 15.25 | .004 |
| R | Superior longitudinal fasciculus | 36 | 11 | 20 | 2 | 22.20 | .004 |
| R | Superior longitudinal fasciculus | 37 | 15 | 17 | 1 | 21.70 | .004 |
| R | Inferior parietal lobe | 44 | -50 | 35 | 1 | 16.64 | .004 |
| R | Superior longitudinal fasciculus | 45 | 4 | 20 | 1 | 25.58 | .004 |
| R | Inferior parietal lobe | 49 | -54 | 16 | 1 | 6.18 | .004 |
| R | Middle temporal lobe | 50 | -14 | -17 | 2 | 11.40 | .004 |
| R | Superior longitudinal fasciculus | 52 | -30 | -10 | 1 | 16.51 | .004 |
| L | Superior longitudinal fasciculus | -42 | -46 | 7 | 1 | 18.45 | .005 |
| L | Pontine crossing tract | -2 | -27 | -27 | 1 | 21.37 | .005 |
| R | Cerebellum | 11 | -48 | -19 | 1 | 12.21 | .005 |
| R | Orbitofrontal area | 14 | 15 | -11 | 1 | 12.54 | .005 |
| R | Optic radiation | 20 | -92 | 4 | 1 | 16.57 | .005 |
| R | Inferior longitudinal fasciculus | 41 | -71 | 8 | 1 | 10.70 | .005 |
| R | Superior longitudinal fasciculus | 41 | -56 | 34 | 1 | 9.57 | .005 |
| R | Inferior longitudinal fasciculus | 45 | -6 | -16 | 1 | 8.78 | .005 |
| *Note.* Region refers to area of highest *t*-value. L = left, R = right. | | | | | | | |

| **Table S3. Between-group contrast of the difference between Scan 2 and Scan 1, Double Trouble > Self-Ordered Search, full list of significant regions** | | | | | | | |
| --- | --- | --- | --- | --- | --- | --- | --- |
| **Region** | | **MNI coordinates** | | | **Size (voxels)** | ***t*-value** | ***p*-value** |
|  |  | **x** | **y** | **z** |  |  |  |
| L | Superior longitudinal fasciculus | -53 | -17 | -18 | 1 | 22.49 | .001 |
| L | Superior temporal lobe | -52 | -36 | 10 | 13 | 27.02 | .001 |
| L | Superior longitudinal fasciculus | -49 | -35 | -12 | 6 | 37.18 | .001 |
| L | Inferior longitudinal fasciculus | -46 | -29 | -12 | 22 | 38.56 | .001 |
| L | Inferior longitudinal fasciculus | -46 | -7 | -15 | 2 | 22.34 | .001 |
| L | Superior longitudinal fasciculus | -44 | -50 | 31 | 2 | 19.37 | .001 |
| L | Inferior occipitofrontal fasciculus | -42 | 31 | 3 | 2 | 24.18 | .001 |
| L | Inferior longitudinal fasciculus | -41 | -8 | -26 | 44 | 35.79 | .001 |
| L | Inferior longitudinal fasciculus | -40 | -4 | -35 | 17 | 36.38 | .001 |
| L | Inferior longitudinal fasciculus | -39 | 4 | -30 | 21 | 28.62 | .001 |
| L | Occipital fusiform area | -39 | -56 | -14 | 7 | 21.81 | .001 |
| L | Inferior occipitofrontal fasciculus | -38 | -11 | -13 | 144 | 63.41 | .001 |
| L | Superior longitudinal fasciculus | -37 | 9 | 17 | 17 | 20.97 | .001 |
| L | Uncinate fasciculus | -35 | 17 | 18 | 61 | 48.43 | .001 |
| L | Inferior longitudinal fasciculus | -35 | -5 | -33 | 7 | 26.47 | .001 |
| L | External capsule | -34 | -7 | 3 | 4 | 40.69 | .001 |
| L | Uncinate fasciculus | -33 | 46 | -1 | 12 | 18.24 | .001 |
| L | Anterior thalamic radiation | -30 | 31 | 22 | 55 | 28.36 | .001 |
| L | Optic radiation | -30 | -21 | -6 | 3 | 30.05 | .001 |
| L | External capsule | -28 | 17 | 0 | 36 | 35.05 | .001 |
| L | External capsule | -28 | 12 | 6 | 19 | 13.23 | .001 |
| L | Middle frontal lobe | -27 | 24 | 31 | 13 | 24.79 | .001 |
| L | Optic radiation | -26 | -27 | -7 | 4 | 26.96 | .001 |
| L | Premotor area | -25 | -1 | 44 | 2 | 12.70 | .001 |
| L | Uncinate fasciculus | -25 | 38 | 17 | 1 | 17.63 | .001 |
| L | Orbitofrontal area | -24 | 23 | -13 | 13 | 28.42 | .001 |
| L | Orbitofrontal area | -23 | 10 | -19 | 10 | 27.86 | .001 |
| L | Fornix | -23 | -41 | 5 | 2 | 12.12 | .001 |
| L | Forceps minor | -18 | 44 | 6 | 12 | 37.05 | .001 |
| L | Anterior thalamic radiation | -17 | -29 | 13 | 22 | 36.03 | .001 |
| L | Callosal body | -17 | 33 | -9 | 5 | 44.48 | .001 |
| L | Anterior thalamic radiation | -16 | -25 | 16 | 2 | 15.46 | .001 |
| L | Anterior thalamic radiation | -16 | 17 | -3 | 2 | 15.32 | .001 |
| L | Forceps minor | -15 | 42 | -13 | 20 | 45.17 | .001 |
| L | Forceps minor | -14 | 35 | -13 | 35 | 46.26 | .001 |
| L | Forceps minor | -14 | 54 | -5 | 8 | 27.75 | .001 |
| L | Anterior thalamic radiation | -9 | -18 | 16 | 1 | 21.48 | .001 |
| L | Cingulum | -9 | 35 | -5 | 1 | 10.40 | .001 |
| L | Cingulum | -8 | -47 | 28 | 3 | 31.93 | .001 |
| R | Anterior thalamic radiation | 2 | -13 | -7 | 14 | 24.26 | .001 |
| R | Pontine crossing tract | 2 | -31 | -37 | 2 | 17.36 | .001 |
| R | Medial frontal lobe | 7 | 46 | -19 | 1 | 7.77 | .001 |
| R | Posterior limb of internal capsule | 15 | -4 | 2 | 21 | 49.59 | .001 |
| R | Visual area V1 | 17 | -96 | 6 | 2 | 18.29 | .001 |
| R | Callosal body | 21 | 26 | 22 | 2 | 10.85 | .001 |
| R | Cerebellum | 22 | -54 | -33 | 5 | 27.27 | .001 |
| R | Optic radiation | 23 | -83 | 5 | 29 | 21.18 | .001 |
| R | External capsule | 30 | 14 | -2 | 11 | 30.46 | .001 |
| R | Superior longitudinal fasciculus | 37 | -50 | 27 | 8 | 25.52 | .001 |
| R | Inferior parietal lobe | 37 | -68 | 28 | 6 | 24.22 | .001 |
| R | Cerebellum | 37 | -68 | -31 | 6 | 41.02 | .001 |
| R | Superior longitudinal fasciculus | 38 | -53 | 21 | 7 | 24.72 | .001 |
| R | Inferior occipitofrontal fasciculus | 39 | 10 | -9 | 17 | 16.28 | .001 |
| R | Cerebellum | 39 | -58 | -36 | 5 | 23.30 | .001 |
| R | Inferior longitudinal fasciculus | 45 | 10 | -21 | 5 | 25.88 | .001 |
| R | Inferior longitudinal fasciculus | 45 | 4 | -32 | 1 | 15.56 | .001 |
| R | Inferior longitudinal fasciculus | 46 | -7 | -14 | 1 | 12.18 | .001 |
| R | Premotor area | 49 | 4 | 31 | 8 | 34.41 | .001 |
| R | Inferior longitudinal fasciculus | 50 | -36 | -14 | 2 | 9.90 | .001 |
| R | Inferior parietal lobe | 53 | -39 | 33 | 8 | 20.73 | .001 |
| R | Primary somatosensory area | 55 | -10 | 32 | 13 | 23.66 | .001 |
| L | Planum temporale | -57 | -19 | 1 | 7 | 18.15 | .002 |
| L | Primary somatosensory area | -53 | -13 | 33 | 7 | 20.96 | .002 |
| L | Middle temporal lobe | -53 | -7 | -24 | 2 | 16.63 | .002 |
| L | Middle temporal lobe | -51 | -10 | -23 | 3 | 18.86 | .002 |
| L | Superior longitudinal fasciculus | -48 | -25 | 29 | 2 | 28.92 | .002 |
| L | Inferior longitudinal fasciculus | -43 | -4 | -23 | 13 | 29.68 | .002 |
| L | Acoustic radiation | -42 | -29 | 3 | 2 | 22.10 | .002 |
| L | External capsule | -34 | -1 | -2 | 16 | 21.39 | .002 |
| L | Acoustic radiation | -34 | -22 | -3 | 7 | 40.81 | .002 |
| L | Optic radiation | -33 | -63 | 13 | 2 | 12.04 | .002 |
| L | Optic radiation | -28 | -19 | -7 | 9 | 34.63 | .002 |
| L | Anterior thalamic radiation | -25 | 28 | 20 | 8 | 27.70 | .002 |
| L | Anterior thalamic radiation | -25 | 29 | 23 | 1 | 5.65 | .002 |
| L | Hippocampus | -24 | -17 | -8 | 1 | 15.11 | .002 |
| L | Primary motor area | -20 | -32 | 52 | 8 | 42.43 | .002 |
| L | Optic radiation | -18 | -93 | 8 | 1 | 9.67 | .002 |
| L | Forceps minor | -17 | 53 | 2 | 5 | 27.43 | .002 |
| L | Forceps minor | -14 | 53 | -7 | 2 | 24.82 | .002 |
| L | Visual area V1 | -12 | -70 | 3 | 1 | 12.23 | .002 |
| L | Visual area V1 | -11 | -76 | 19 | 1 | 17.23 | .002 |
| L | Cingulum | -11 | -22 | 35 | 1 | 7.41 | .002 |
| L | Anterior thalamic radiation | -9 | -20 | 13 | 1 | 15.28 | .002 |
| L | Anterior thalamic radiation | -9 | 6 | -7 | 1 | 22.73 | .002 |
| R | Anterior thalamic radiation | 6 | -20 | -13 | 2 | 17.55 | .002 |
| R | Anterior thalamic radiation | 6 | -11 | 12 | 1 | 15.42 | .002 |
| R | Forceps minor | 12 | 53 | 30 | 3 | 21.34 | .002 |
| R | Anterior thalamic radiation | 13 | -1 | 1 | 2 | 39.38 | .002 |
| R | Corticospinal tract | 15 | -41 | 66 | 2 | 15.93 | .002 |
| R | Forceps major | 16 | -90 | 1 | 9 | 14.12 | .002 |
| R | Putamen | 18 | 5 | -7 | 1 | 19.49 | .002 |
| R | Superior longitudinal fasciculus | 23 | -59 | 51 | 6 | 10.81 | .002 |
| R | Optic radiation | 25 | -91 | 1 | 3 | 13.98 | .002 |
| R | Inferior occipitofrontal fasciculus | 25 | -71 | 30 | 1 | 22.63 | .002 |
| R | Superior longitudinal fasciculus | 26 | 10 | 43 | 1 | 14.98 | .002 |
| R | Cerebellum | 32 | -51 | -38 | 2 | 19.75 | .002 |
| R | Inferior longitudinal fasciculus | 34 | -67 | -8 | 2 | 10.92 | .002 |
| R | Inferior parietal lobe | 37 | -71 | 34 | 3 | 17.63 | .002 |
| R | Inferior longitudinal fasciculus | 38 | -36 | -18 | 2 | 17.53 | .002 |
| R | Temporal occipital fusiform area | 39 | -54 | -13 | 1 | 21.15 | .002 |
| R | Corticospinal tract | 40 | -5 | 33 | 7 | 33.64 | .002 |
| R | Superior longitudinal fasciculus | 47 | -40 | 31 | 7 | 23.47 | .002 |
| R | Middle temporal gyrus | 57 | -21 | -14 | 9 | 16.89 | .002 |
| R | Planum temporale | 60 | -19 | 7 | 1 | 15.81 | .002 |
| L | Superior longitudinal fasciculus | -51 | -12 | -22 | 1 | 10.91 | .003 |
| L | Superior longitudinal fasciculus | -46 | -47 | 0 | 1 | 17.93 | .003 |
| L | Inferior longitudinal fasciculus | -41 | -3 | -30 | 6 | 22.68 | .003 |
| L | Middle frontal lobe | -37 | 33 | 24 | 1 | 14.29 | .003 |
| L | Inferior longitudinal fasciculus | -33 | 3 | -31 | 2 | 23.50 | .003 |
| L | External capsule | -32 | -8 | 8 | 5 | 13.00 | .003 |
| L | Retrolenticular internal capsule | -31 | -21 | -2 | 4 | 44.97 | .003 |
| L | External capsule | -31 | 8 | 5 | 1 | 17.06 | .003 |
| L | External capsule | -29 | 9 | 7 | 1 | 20.76 | .003 |
| L | Superior frontal lobe | -24 | 3 | 44 | 2 | 15.64 | .003 |
| L | Anterior thalamic radiation | -23 | -32 | 10 | 4 | 22.38 | .003 |
| L | Callosal body | -22 | 36 | 5 | 4 | 36.02 | .003 |
| L | Hippocampus | -20 | -39 | 7 | 3 | 26.33 | .003 |
| L | Superior occipitofrontal fasciculus | -19 | -2 | 26 | 1 | 14.23 | .003 |
| L | Callosal body | -17 | 38 | -3 | 6 | 55.98 | .003 |
| L | Callosal body | -14 | 35 | -2 | 5 | 55.83 | .003 |
| L | Anterior thalamic radiation | -14 | -22 | 16 | 3 | 14.87 | .003 |
| L | Optic radiation | -14 | -77 | 2 | 2 | 14.38 | .003 |
| L | Cingulum | -13 | -40 | 40 | 1 | 18.60 | .003 |
| L | Corticospinal tract | -10 | -34 | 56 | 4 | 22.40 | .003 |
| L | Accumbens | -6 | 7 | -7 | 4 | 12.57 | .003 |
| R | Anterior thalamic radiation | 2 | -18 | -10 | 2 | 24.82 | .003 |
| R | Anterior thalamic radiation | 8 | 1 | 6 | 1 | 13.20 | .003 |
| R | Cingulum | 12 | -50 | 19 | 5 | 17.18 | .003 |
| R | Inferior occipitofrontal fasciculus | 25 | 30 | 11 | 6 | 34.29 | .003 |
| R | Inferior longitudinal fasciculus | 37 | -77 | 15 | 2 | 22.40 | .003 |
| R | Superior longitudinal fasciculus | 39 | -48 | 26 | 3 | 13.44 | .003 |
| R | Inferior occipitofrontal fasciculus | 42 | 35 | 2 | 1 | 7.65 | .003 |
| R | Inferior occipitofrontal fasciculus | 44 | 31 | 8 | 3 | 18.34 | .003 |
| R | Visual area V5 | 44 | -68 | 10 | 2 | 10.98 | .003 |
| R | Inferior occipitofrontal fasciculus | 46 | 30 | 5 | 4 | 11.46 | .003 |
| R | Primary somatosensory area | 51 | -12 | 32 | 2 | 19.95 | .003 |
| L | Superior longitudinal fasciculus | -55 | -6 | 17 | 1 | 13.23 | .004 |
| L | Inferior longitudinal fasciculus | -46 | 1 | -19 | 1 | 16.05 | .004 |
| L | Inferior occipitofrontal fasciculus | -42 | 31 | 1 | 3 | 24.30 | .004 |
| L | Inferior longitudinal fasciculus | -38 | -5 | -31 | 4 | 25.85 | .004 |
| L | Temporal occipital fusiform area | -38 | -55 | -12 | 1 | 17.75 | .004 |
| L | Insula | -36 | 11 | -10 | 1 | 8.87 | .004 |
| L | Optic radiation | -35 | -63 | 15 | 1 | 7.53 | .004 |
| L | Premotor area | -34 | -11 | 54 | 1 | 13.61 | .004 |
| L | Inferior longitudinal fasciculus | -33 | -2 | -29 | 4 | 33.77 | .004 |
| L | External capsule | -33 | 5 | 0 | 3 | 23.87 | .004 |
| L | External capsule | -33 | 0 | 2 | 2 | 31.04 | .004 |
| L | Uncinate fasciculus | -33 | 46 | -4 | 1 | 14.66 | .004 |
| L | Corticospinal tract | -31 | -17 | 34 | 1 | 14.04 | .004 |
| L | External capsule | -30 | -7 | 15 | 3 | 21.52 | .004 |
| L | Corticospinal tract | -27 | -9 | 19 | 4 | 41.83 | .004 |
| L | Anterior thalamic radiation | -26 | 34 | 13 | 2 | 24.66 | .004 |
| L | Anterior thalamic radiation | -26 | 34 | 11 | 1 | 20.87 | .004 |
| L | Orbitofrontal area | -24 | 19 | -15 | 2 | 19.60 | .004 |
| L | Anterior thalamic radiation | -13 | -19 | 8 | 1 | 9.29 | .004 |
| L | Anterior corona radiata | -12 | 21 | -11 | 2 | 14.85 | .004 |
| L | Callosal body | -12 | 23 | -9 | 1 | 13.83 | .004 |
| L | Anterior thalamic radiation | -4 | -11 | -4 | 1 | 7.18 | .004 |
| L | Anterior thalamic radiation | -3 | -11 | -8 | 3 | 9.94 | .004 |
| L | Fornix | -1 | -9 | 9 | 1 | 6.95 | .004 |
| R | Fornix | 2 | 5 | -5 | 1 | 16.85 | .004 |
| R | Fornix | 4 | 1 | 1 | 1 | 19.02 | .004 |
| R | Fornix | 6 | 5 | -5 | 2 | 13.78 | .004 |
| R | Anterior thalamic radiation | 6 | -5 | 7 | 2 | 13.72 | .004 |
| R | Fornix | 9 | -29 | 16 | 1 | 14.99 | .004 |
| R | Corticospinal tract | 12 | -25 | -6 | 2 | 20.82 | .004 |
| R | Forceps major | 16 | -83 | 29 | 5 | 9.64 | .004 |
| R | Pallidum | 16 | 7 | -4 | 1 | 4.21 | .004 |
| R | Forceps minor | 19 | 43 | -1 | 4 | 26.57 | .004 |
| R | Corticospinal tract | 27 | -15 | 27 | 1 | 24.62 | .004 |
| R | Cerebellum | 29 | -66 | -41 | 1 | 7.84 | .004 |
| R | Optic radiation | 38 | -51 | 23 | 3 | 19.49 | .004 |
| R | Visual area V5 | 41 | -64 | 11 | 1 | 6.89 | .004 |
| R | Corticospinal tract | 42 | -4 | 41 | 1 | 16.93 | .004 |
| R | Superior longitudinal fasciculus | 43 | -38 | 32 | 2 | 18.81 | .004 |
| R | Superior longitudinal fasciculus | 44 | -38 | 42 | 1 | 13.74 | .004 |
| R | Inferior occipitofrontal fasciculus | 45 | 29 | 7 | 1 | 17.36 | .004 |
| R | Parietal operculum | 58 | -7 | 15 | 1 | 10.23 | .004 |
| L | Inferior longitudinal fasciculus | -45 | -25 | -15 | 2 | 25.44 | .005 |
| L | Inferior longitudinal fasciculus | -41 | -4 | -28 | 1 | 21.05 | .005 |
| L | Inferior longitudinal fasciculus | -40 | -6 | -27 | 1 | 25.91 | .005 |
| L | Inferior parietal lobe | -36 | -57 | 19 | 1 | 8.62 | .005 |
| L | Orbitofrontal area | -35 | 26 | -10 | 1 | 12.92 | .005 |
| L | Inferior occipitofrontal fasciculus | -34 | 2 | -9 | 1 | 11.36 | .005 |
| L | Retrolenticular internal capsule | -31 | -32 | 8 | 1 | 35.59 | .005 |
| L | Fornix | -30 | -29 | -3 | 1 | 26.54 | .005 |
| L | Orbitofrontal area | -30 | 21 | -18 | 1 | 16.10 | .005 |
| L | Uncinate fasciculus | -29 | 5 | -10 | 1 | 15.49 | .005 |
| L | Corticospinal tract | -27 | -11 | 18 | 1 | 45.98 | .005 |
| L | Orbitofrontal area | -27 | 19 | -19 | 1 | 14.90 | .005 |
| L | Corticospinal tract | -26 | -7 | 31 | 1 | 22.82 | .005 |
| L | Callosal body | -22 | -52 | 11 | 2 | 28.18 | .005 |
| L | Splenium of corpus callosum | -21 | -52 | 17 | 2 | 25.43 | .005 |
| L | Callosal body | -21 | 34 | 11 | 1 | 26.98 | .005 |
| L | Corticospinal tract | -21 | -3 | 17 | 1 | 27.24 | .005 |
| L | Superior parietal lobe | -20 | -39 | 39 | 1 | 25.26 | .005 |
| L | Splenium of corpus callosum | -18 | -49 | 16 | 1 | 30.45 | .005 |
| L | Anterior thalamic radiation | -13 | -24 | 16 | 1 | 13.88 | .005 |
| L | Superior parietal lobe | -10 | -51 | 51 | 1 | 9.45 | .005 |
| L | Fornix | -1 | -15 | 19 | 1 | 11.22 | .005 |
| R | Anterior thalamic radiation | 5 | -2 | 6 | 1 | 8.67 | .005 |
| R | Forceps minor | 13 | 51 | 28 | 1 | 11.66 | .005 |
| R | Forceps major | 13 | -84 | 34 | 1 | 9.93 | .005 |
| R | Anterior thalamic radiation | 16 | -14 | 21 | 1 | 8.67 | .005 |
| R | Anterior thalamic radiation | 17 | -12 | 19 | 1 | 8.03 | .005 |
| R | Cingulum | 17 | -55 | 36 | 1 | 20.53 | .005 |
| R | Callosal body | 18 | 28 | 28 | 1 | 23.24 | .005 |
| R | Callosal body | 21 | 21 | 26 | 1 | 13.27 | .005 |
| R | Cerebellum | 21 | -59 | -32 | 1 | 26.19 | .005 |
| R | Superior parietal lobe | 22 | -61 | 51 | 1 | 8.23 | .005 |
| R | Corticospinal tract | 26 | 4 | 21 | 2 | 24.17 | .005 |
| R | Superior longitudinal fasciculus | 31 | -43 | 43 | 1 | 11.89 | .005 |
| R | Inferior occipitofrontal fasciculus | 37 | -71 | 18 | 1 | 11.94 | .005 |
| R | Superior longitudinal fasciculus | 37 | -52 | 26 | 1 | 9.28 | .005 |
| R | Superior longitudinal fasciculus | 39 | -51 | 28 | 1 | 12.65 | .005 |
| R | Superior longitudinal fasciculus | 42 | -37 | 30 | 1 | 18.07 | .005 |
| R | Planum temporale | 54 | -33 | 12 | 1 | 22.06 | .005 |
| R | Parietal operculum | 57 | -5 | 15 | 1 | 9.05 | .005 |
| R | Superior temporal lobe | 61 | -25 | 7 | 1 | 18.79 | .005 |
| R | Superior temporal lobe | 62 | -27 | 7 | 1 | 14.20 | .005 |
| *Note.* Region refers to area of highest *t*-value. L = left, R = right. | | | | | | | |

| **Table S4. Between-group contrast of the difference between Scan 2 and Scan 1, Self-Ordered Search > Double Trouble, full list of significant regions** | | | | | | | |
| --- | --- | --- | --- | --- | --- | --- | --- |
| **Region** | | **MNI coordinates** | | | **Size (voxels)** | ***t*-value** | ***p*-value** |
|  |  | **x** | **y** | **z** |  |  |  |
| L | Superior longitudinal fasciculus | -52 | -41 | -6 | 4 | 30.68 | .001 |
| L | Acoustic radiation | -52 | -6 | -3 | 3 | 21.25 | .001 |
| L | Superior longitudinal fasciculus | -45 | -45 | 25 | 4 | 18.27 | .001 |
| L | Inferior longitudinal fasciculus | -37 | -81 | 14 | 2 | 13.58 | .001 |
| L | Anterior thalamic radiation | -33 | 28 | 27 | 11 | 36.56 | .001 |
| L | Visual area V4 | -32 | -80 | -4 | 13 | 20.30 | .001 |
| L | Optic radiation | -20 | -79 | 22 | 10 | 32.30 | .001 |
| R | Primary motor area | 7 | -28 | 53 | 10 | 24.34 | .001 |
| R | Precuneus | 7 | -54 | 14 | 9 | 26.45 | .001 |
| R | Premotor area | 13 | -17 | 61 | 5 | 29.81 | .001 |
| R | Primary motor area | 15 | -36 | 68 | 5 | 32.06 | .001 |
| R | Cerebellum | 22 | -41 | -32 | 2 | 15.30 | .001 |
| R | Superior parietal lobe | 24 | -59 | 47 | 10 | 20.22 | .001 |
| R | Cerebellum | 24 | -52 | -41 | 6 | 15.71 | .001 |
| R | Inferior occipitofrontal fasciculus | 37 | 15 | -5 | 6 | 26.74 | .001 |
| R | Inferior longitudinal fasciculus | 40 | -5 | -30 | 2 | 15.69 | .001 |
| R | Superior longitudinal fasciculus | 40 | 0 | 31 | 1 | 22.99 | .001 |
| R | Inferior longitudinal fasciculus | 42 | -29 | -12 | 10 | 46.58 | .001 |
| R | Corticospinal tract | 48 | -1 | 35 | 3 | 15.56 | .001 |
| R | Inferior frontal lobe | 48 | 17 | 18 | 2 | 26.99 | .001 |
| R | Superior longitudinal fasciculus | 51 | -47 | 3 | 2 | 17.76 | .001 |
| R | Superior longitudinal fasciculus | 53 | -16 | -17 | 1 | 13.26 | .001 |
| L | Inferior longitudinal fasciculus | -33 | -56 | -11 | 2 | 16.10 | .002 |
| L | Superior parietal lobe | -23 | -59 | 46 | 2 | 22.32 | .002 |
| R | Anterior thalamic radiation | 19 | -29 | 7 | 5 | 11.39 | .002 |
| R | Anterior thalamic radiation | 20 | 21 | 2 | 11 | 33.14 | .002 |
| R | Anterior thalamic radiation | 25 | -74 | 26 | 2 | 31.12 | .002 |
| R | Inferior occipitofrontal fasciculus | 29 | 36 | 3 | 1 | 20.34 | .002 |
| R | Optic radiation | 34 | -60 | 0 | 4 | 30.01 | .002 |
| R | Inferior parietal lobe | 37 | -80 | 22 | 5 | 19.29 | .002 |
| R | Inferior occipitofrontal fasciculus | 37 | 35 | 7 | 1 | 14.12 | .002 |
| R | Superior longitudinal fasciculus | 40 | -2 | 36 | 4 | 23.10 | .002 |
| R | Inferior longitudinal fasciculus | 40 | -41 | -11 | 2 | 12.10 | .002 |
| R | Superior longitudinal fasciculus | 43 | 11 | 13 | 8 | 20.79 | .002 |
| R | Optic radiation | 43 | -42 | -1 | 5 | 18.20 | .002 |
| R | Superior longitudinal fasciculus | 45 | -44 | -1 | 4 | 24.10 | .002 |
| R | Superior longitudinal fasciculus | 48 | -44 | -2 | 9 | 31.04 | .002 |
| L | Superior longitudinal fasciculus | -51 | -32 | -10 | 2 | 16.03 | .003 |
| L | Inferior longitudinal fasciculus | -49 | -3 | -27 | 1 | 15.03 | .003 |
| L | Superior longitudinal fasciculus | -48 | -44 | -2 | 3 | 20.81 | .003 |
| L | Inferior longitudinal fasciculus | -34 | -71 | 21 | 1 | 21.06 | .003 |
| L | Optic radiation | -33 | -78 | 0 | 3 | 18.07 | .003 |
| L | Amygdala | -25 | -9 | -8 | 1 | 10.17 | .003 |
| L | Cingulum | -9 | -13 | 33 | 3 | 23.31 | .003 |
| L | Cingulum | -8 | -17 | 34 | 3 | 19.33 | .003 |
| L | Visual area V2 | -8 | -83 | 22 | 1 | 12.94 | .003 |
| R | Optic radiation | 13 | -84 | -1 | 3 | 15.89 | .003 |
| R | Cerebellum | 17 | -76 | -28 | 5 | 24.33 | .003 |
| R | Hippocampus | 17 | -35 | -3 | 1 | 8.69 | .003 |
| R | Callosal body | 19 | -49 | 37 | 1 | 8.99 | .003 |
| R | Cerebellum | 27 | -56 | -37 | 10 | 40.72 | .003 |
| R | Middle frontal lobe | 29 | 31 | 33 | 1 | 9.64 | .003 |
| R | Inferior occipitofrontal fasciculus | 31 | 18 | 20 | 1 | 6.73 | .003 |
| R | Inferior occipitofrontal fasciculus | 33 | 39 | 1 | 1 | 38.25 | .003 |
| R | Visual area V5 | 39 | -68 | 10 | 1 | 11.95 | .003 |
| R | Inferior parietal lobe | 42 | -66 | 26 | 5 | 13.43 | .003 |
| R | Superior longitudinal fasciculus | 47 | -51 | -1 | 1 | 10.18 | .003 |
| L | Corticospinal tract | -33 | -10 | 41 | 3 | 46.65 | .004 |
| L | Corticospinal tract | -29 | -19 | 49 | 1 | 33.12 | .004 |
| L | Uncinate fasciculus | -27 | 27 | -10 | 1 | 18.05 | .004 |
| L | Primary somatosensory area | -22 | -35 | 64 | 2 | 20.82 | .004 |
| L | Anterior thalamic radiation | -14 | -16 | 18 | 1 | 9.04 | .004 |
| L | Optic radiation | -10 | -84 | 21 | 1 | 14.14 | .004 |
| R | Visual area V2 | 8 | -88 | 19 | 1 | 20.65 | .004 |
| R | Cerebellum | 13 | -73 | -30 | 1 | 20.14 | .004 |
| R | Callosal body | 24 | -45 | 2 | 1 | 11.47 | .004 |
| R | Amygdala | 24 | -8 | -4 | 1 | 5.13 | .004 |
| R | Anterior thalamic radiation | 28 | 33 | 18 | 1 | 5.89 | .004 |
| R | Primary somatosensory area | 28 | -34 | 63 | 1 | 12.75 | .004 |
| R | Optic radiation | 31 | -64 | 15 | 1 | 7.20 | .004 |
| R | Anterior thalamic radiation | 33 | 50 | 8 | 1 | 7.72 | .004 |
| R | Inferior occipitofrontal fasciculus | 33 | 41 | 2 | 1 | 24.45 | .004 |
| R | Hippocampus | 36 | -30 | -16 | 1 | 5.38 | .004 |
| R | Inferior longitudinal fasciculus | 45 | -11 | -19 | 3 | 4.90 | .004 |
| L | Planum temporale | -57 | -28 | 9 | 1 | 11.59 | .005 |
| L | Primary motor cortex | -36 | -31 | 50 | 1 | 25.99 | .005 |
| L | Corticospinal tract | -35 | -9 | 41 | 1 | 33.92 | .005 |
| L | Superior parietal lobe | -21 | -52 | 58 | 1 | 7.75 | .005 |
| L | Cerebellum | -11 | -48 | -19 | 1 | 12.10 | .005 |
| L | Visual area V2 | -8 | -89 | 17 | 1 | 13.95 | .005 |
| L | Cingulum | -8 | -6 | 36 | 1 | 17.95 | .005 |
| R | Primary motor area | 6 | -22 | 52 | 2 | 14.37 | .005 |
| R | Visual area V2 | 8 | -92 | 17 | 1 | 16.09 | .005 |
| R | Visual area V2 | 13 | -72 | 3 | 1 | 9.43 | .005 |
| R | Cingulum | 23 | -42 | -3 | 1 | 23.33 | .005 |
| R | Uncinate fasciculus | 24 | 7 | -13 | 1 | 16.03 | .005 |
| R | Visual area V3 | 25 | -85 | -6 | 1 | 11.75 | .005 |
| R | Visual area V2 | 26 | -91 | 12 | 1 | 12.61 | .005 |
| R | Anterior thalamic radiation | 32 | 36 | 16 | 1 | 16.38 | .005 |
| R | Premotor area | 38 | -2 | 38 | 2 | 34.31 | .005 |
| R | Inferior occipitofrontal fasciculus | 38 | 38 | -3 | 1 | 14.55 | .005 |
| R | Temporal fusiform area | 40 | -5 | -32 | 1 | 5.35 | .005 |
| R | Superior longitudinal fasciculus | 42 | 0 | 32 | 1 | 22.00 | .005 |
| R | Superior longitudinal fasciculus | 45 | 1 | 22 | 2 | 31.65 | .005 |
| R | Superior longitudinal fasciculus | 46 | 2 | 20 | 1 | 26.01 | .005 |
| R | Middle temporal lobe | 57 | -14 | -18 | 1 | 19.83 | .005 |
| *Note.* Region refers to area of highest *t*-value. L = left, R = right. | | | | | | | |
